# An atlas of protein-protein associations of human tissues prioritizes candidate disease genes

**DOI:** 10.1101/2024.05.15.594301

**Authors:** Diederik S Laman Trip, Marc van Oostrum, Danish Memon, Fabian Frommelt, Delora Baptista, Kalpana Panneerselvam, Glyn Bradley, Luana Licata, Henning Hermjakob, Sandra Orchard, Gosia Trynka, Ellen McDonagh, Andrea Fossati, Ruedi Aebersold, Matthias Gstaiger, Bernd Wollscheid, Pedro Beltrao

## Abstract

Proteins that interact together participate in the same cellular process and influence the same organismal traits. Despite the progress in mapping protein-protein interactions we lack knowledge of how they differ between tissues. Due to coordinated (post)transcriptional control, protein complex members have highly correlated abundances that are predictive of functional association. Here, we have compiled 7873 proteomic samples measuring protein levels in 11 human tissues and use these to define an atlas with tissue-specific protein associations. This method recapitulates known protein complexes and the larger structural organization of the cell. Interactions of stable protein complexes are well preserved across tissues, while signaling and metabolic interactions show larger variation. Further, we find that less than 18% of differences between tissues are estimated to be due to differences in gene expression while cell-type specific cellular structures, such as synaptic components, represent a significant driver of differences between tissues. We further supported the brain protein association network through co-fractionation experiments in synaptosomes, curation of brain derived pull-down data and AlphaFold2 models. Together these results illustrate how this brain specific protein interaction network can functionally prioritize candidate genes within loci linked to brain disorders.

## INTRODUCTION

Protein-protein interactions mediate the biophysical structure and functioning of the cell, with disruptions potentially leading to disease. Unraveling the universe of interactions for human proteins has been a long-standing challenge [1], with recent advances in mass spectrometry coupled with high-throughput screens such as protein-fragment complementation [2], yeast two-hybrid (Y2H) [3], affinity-purification (AP-MS) [4,5], or co-fractionation (CF-MS) [6–8] enabling the discovery of interactions on a proteome-wide scale. These experimental datasets are complemented by modern computational approaches that compile collections of interactions using machine learning-based methods often trained with experimental evidence [1]. Together, these experimental and computational efforts are the basis for a myriad of publicly available, high-quality protein interaction databases [1,9–11].

However, these databases typically aggregate interactions without specifying context, while the interactome is highly tissue or cell-state specific with less than half of the proteome being detected in all tissues [12]. Elucidating the tissue-specificity of protein interactions is important for understanding cell-type specific function, finding drug targets, and developing a systems level understanding of the human cell. Indeed, specifying tissue-specificity of the interactome has proven difficult. Initial attempts to generate tissue specific interaction networks have relied on gene expression data to identify co-expressed genes or to exclude proteins based on the lack of expression in a given tissue [13,14]. However, the accuracy of predicting protein associations by mRNA co-expression is limited and it is unclear to what extent gene expression is the major driver of changes in protein interactions [15–18]. While there are some recent experimental efforts for establishing tissue-specific interactions on a proteome-wide scale [4,6], such works require extensive resources even for a single tissue and often use immortalized cell lines or other models whose interactome may not appropriately represent the human tissue. One approach has used changes in ratios for complex subunits from proteomic measurements to establish changes in protein interactions in a disease state [19]. An alternative approach is to use the co-abundance of proteins for the purpose of establishing protein associations [5,17,20,21]. Co-abundance has been shown to be an accurate measure for predicting protein-protein associations likely due to the fact that protein complexes consist of subunits assembled in defined stoichiometries that are often co-expressed and with orphaned subunits often being degraded [15,22–25]. The accuracy of co-abundance for predicting protein association and a rising number of large-scale proteomics studies of human cancers – with genetic heterogeneity underlying diverse cellular changes – offer a timely opportunity to systematically establish the tissue-specificity of protein associations.

Here, we present a protein association atlas derived from the co-abundance of proteins in 7,840 human biopsies, scoring the likelihood of 114 million protein associations across 11 tissues. This atlas recovers well-known protein complexes, maps relationships between and across traits and cellular components, and proposes context-relevant associations for disease genes. Focusing on the brain and using interactions derived from orthogonal approaches, we illustrate the use of our association atlas for prioritizing genes and interactions related to disease in a tissue-specific manner.

## RESULTS

### Protein co-abundance for scoring protein associations on a genome-wide scale

We started by collecting protein abundance data from proteomics studies of cohorts of cancer patients. In total, we compiled a dataset of 50 studies across 14 human tissues, encompassing 5,696 samples of tumors and 2,177 samples of adjacent healthy tissue (Fig. 1a - Table S1) [26–75]. We further included the mRNA expression data paired to the proteomics for 2,930 of the tumor and 733 of the healthy samples. Following previous studies, we used the fact that protein complex members are strongly transcriptionally and post-transcriptionally co-regulated to compute probabilities of protein-protein associations from the abundance data (Fig. 1b - Methods; Fig. S1) [5,17,20]. In short, we pre-processed the abundance data to obtain a log-transformed and median-normalized abundance across patients. For each study, we then computed a co-abundance estimate of a protein pair as the Pearson correlation when both proteins were quantified in at least 30 samples (Fig. S2). Finally, we converted the co-abundance estimates to association probabilities using a logistic model and curated stable protein complexes as positives (CORUM [76]).

**Figure 1.**
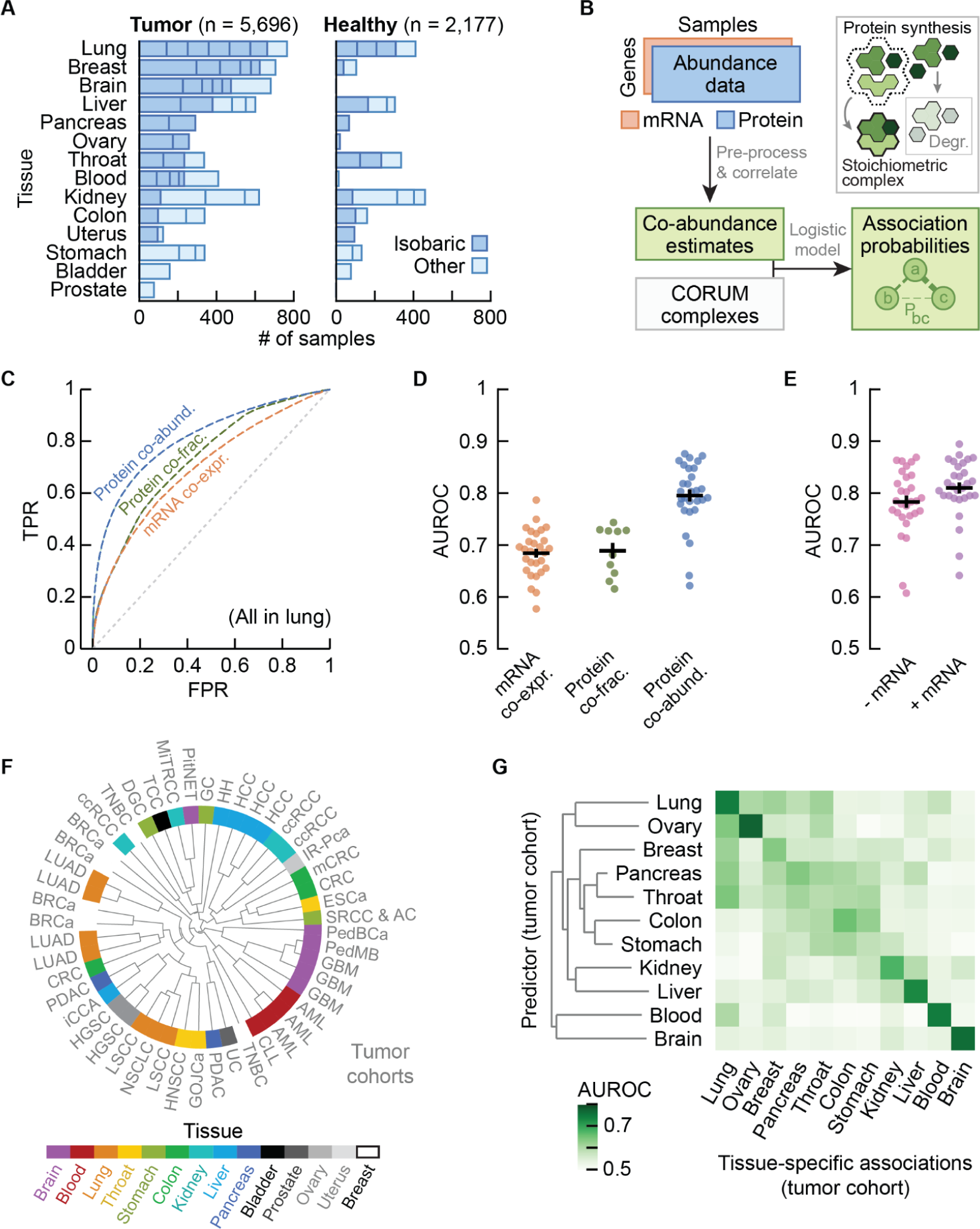
Protein co-abundance outperforms mRNA co-expression and protein co-fractionation for recovering protein-protein interactions on a genome-wide scale. **(a)** Number of tumor and healthy samples per tissue. Bar sections indicate individual studies, with multiplexed proteomics with isobaric labeling (dark blue) or other methods (light blue). **(b)** Schematic representation of workflow. Subunits of protein complexes occur in fixed stoichiometries. Protein or mRNA abundance is used to compute co-abundance estimates, which are converted to protein interaction probabilities using a logistic model with CORUM complexes as positives. **(c)** ROC curves for interaction probabilities in lung tissue derived from protein co-abundance (blue), mRNA co-expression (orange) and protein co-fractionation (green). Grey dotted line shows the performance of a random classifier. **(d)** AUC values for interaction probabilities as illustrated in (c). Shown are studies that quantified both protein co-abundance (blue) and mRNA co-expression (orange), or protein co-fractionation (green). **(e)** AUC values for interaction probabilities derived from protein co-abundance combined with mRNA co-expression through a linear model (purple), and for protein co-abundance after regressing gene expression out of the protein abundance (pink). Shown are the same studies as in (d). In (d-e), each dot represents one study having paired transcriptomics and proteomics data. Protein pairs were filtered for having association probabilities from both modalities. Error bars show mean with s.e.m.. **(f)** Radial dendrogram clustering the 48 cohorts through complete-linkage clustering with the Pearson correlation distance. Cohorts are labeled according to the type of cancer, colors represent the different human tissues. **(g)** Heatmap of AUCs for recovering tissue-specific interactions with cohorts that were withheld when predicting these interactions. Each square represents the average AUC for all cohorts of a given tissue.

To test the ability of the association probabilities to recover known complex members, we computed ROC curves for probabilities derived from protein co-abundance, mRNA co-expression and protein co-fractionation [6,7,77] (Fig. 1c; Fig. S3). We found that protein co-abundance (AUC ∼0.80 ± 0.01) outperformed correlation of protein co-fractionation data (AUC ∼0.69 ± 0.02) and mRNA co-expression (0.68 ± 0.01) for recovering known interactions (Fig. 1d - Methods). In addition, the combination of mRNA and protein abundance data did not significantly improve the recovery of known complex members (Fig. 1e - AUC ∼0.81 ± 0.01, p-value = 0.47; two-sided t-test). We therefore chose to use only protein co-abundance for computing association probabilities, since we don’t observe an improvement and only half of all cohorts had paired mRNA expression data available. Additionally, we found similar AUCs when regressing gene expression out of the protein abundance prior to computing protein co-abundance estimates (∼0.78 ± 0.01, p-value = 0.35), suggesting that post-transcriptional processes – not regulation of gene expression – drive most of the predictive power for protein associations.

Having established that the association probabilities derived from protein co-abundance data recover known interactions of protein complex members, we sought to test whether replicate studies of the same tissue yielded association probabilities that were representative for each tissue. Using the 1,256,313 association probabilities that were quantified for all studies, we found that the replicate cohorts from the same tissue generally clustered together (Fig. 1f – e.g., blood, brain, liver, and lung). Next, we selected the associations that were tissue-specific - associations whose average probability exceeded the 95-th percentile for a given tissue and whose average probability remained below 0.5 across all other tissues. Through a hold-one-out methodology, we found that the tissue-specific associations are primarily recovered by cohorts of the same tissue of origin (AUCs 0.69 ± 0.01) compared to cohorts from different tissues (0.55 ± 0.00 - p-values below 1e-3 for all tissues except the stomach and throat; one sided t-test) (Fig. 1g – Methods; Fig. S4). These observations suggest that the tissue of origin is a major driver of differences between cohorts.

### An atlas of protein associations in human tissues

With the replicate cohorts representing the tissue of origin, we aggregated the replicate association probabilities into single association scores for 11 human tissues (Fig. 2a – Methods; Table S2). Aggregating the replicate cohorts was advantageous, as all but one of the individual cohorts were outperformed by the tissue-level scores for recovering known protein interactions (p-value = 9.4e-4). Moreover, the tumor-derived scores outperformed the healthy-tissue derived scores for all tissues as expected, given that the genetic heterogeneity of tumors increases variation between samples (p-value 6.4e-4 - paired t-test; Fig. S5) and thus yields better co-abundance estimates (Fig. 2b - AUCs 0.87 ± 0.01 and 0.82 ± 0.01, respectively). The healthy- and tumor-derived scores originated from separate dissections of the same tissues and patients, and could thus serve as independent replicates. Analogous to the cohorts, we computed tissue-specific associations from the healthy-tissue derived scores, which we then recovered with the tumor-derived scores (Fig. 2c). For all tissues, we found that the tumor-derived scores primarily recovered the tissue-specific associations of the same healthy tissue (AUC ∼ 0.81 ± 0.2) compared to the other healthy tissues (AUC ∼ 0.53 ± 0.0; p-value = 8.6e-12). These analyses show that the co-abundance derived, tissue-level association scores recover known protein interactions and are reproducible and representative of the tissue of origin (Fig. S6).

**Figure 2.**
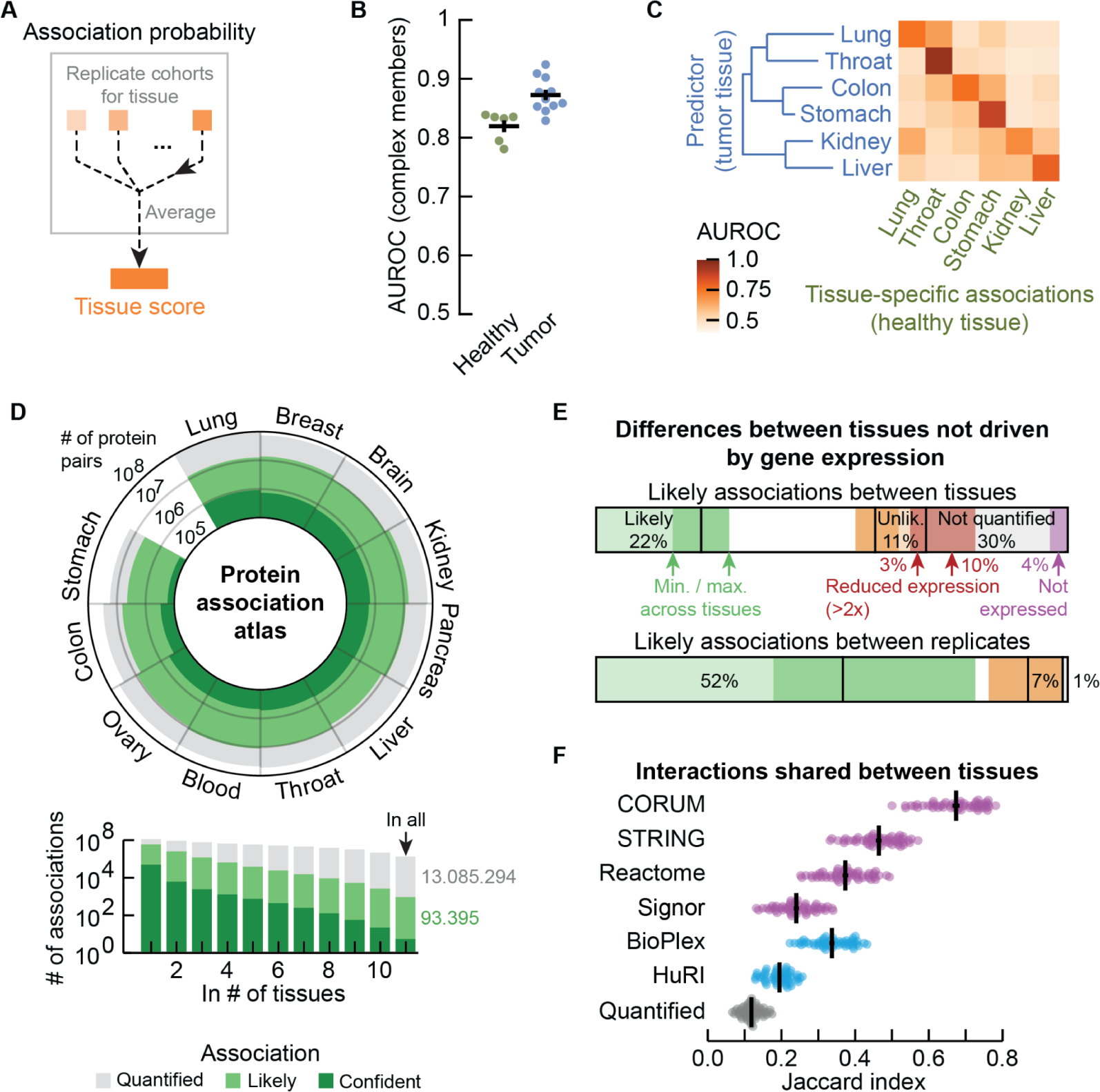
Association atlas scores likelihood of protein interactions across human tissues. **(a)** Schematic for aggregating replicate cohorts into a single tissue score. **(b)** AUC values for the tissue scores derived from healthy samples (green) and tumor samples (blue). Positives defined by complex members in CORUM. Association scores filtered for protein pairs having probabilities in all cohorts of a tissue. **(c)** Heatmap of AUCs for using tumor-derived tissue scores to recover tissue-specific interactions defined by the tissue scores from healthy tissues. Tissue scores only included cohorts that had both healthy and tumor samples. In (b,e) tissues were clustered with average-linkage clustering using the Manhattan distance. **(d)** Atlas of protein associations in 11 human tissues. Radial diagram shows, for each tissue, the number of protein-pairs that were quantified (gray), are likely to interact (light green - interaction score exceeds 0.5) or are confidently quantified (dark green - association score exceeds 99-th percentile of scores in a tissue). Bar graph shows the number of associations that are quantified in a given number of tissues. **(e)** Likely associations of tissues classified in other tissues as likely (also association score > 0.5 - green), unlikely (association score < 0.25 - orange) or not quantified (no association score - gray), and for the latter two categories further subclassified as having reduced expression (>2-fold - red; consensus normalized expression from the Protein Atlas) or not being expressed (purple) in the other tissues. Shown is the percentage of likely associations classified in each category averaged over all pairs of tissues (top bar) and between the replicate tumor- and healthy-derived scores (bottom bar). Dark shaded area shows minimum and maximum percentage for the tissue-level averages. **(f)** Likely associations shared between pairs of tissues as quantified by the Jaccard index (gray dots), compared to shared associations restricted to complex members (CORUM), functional associations (STRING scores exceeding 400), biological pathways (Reactome) and signaling (SIGNOR) (purple dots), or associations detected through yeast two-hybrid (HuRI)) or affinity purification (Bioplex) experiments (blue dots). Each dot represents a pair of tissues. Error bars show mean with s.e.m..

We defined a protein association atlas with association scores for all quantified protein pairs by averaging the association probabilities over the cohorts of each tissue. The resulting association atlas scores the association likelihood for 114 million protein pairs across 11 human tissues (Fig. 2d). On average, each tissue contains association scores for 55 ± 6.1 million protein pairs, of which 10 ± 0.9 million are likely to be associated (score above 0.5), and 0.55 ± 0.06 million are “confident” associations (score exceeds 99-th percentile of the scores in a tissue). Finally, we found that protein associations tend to be likely in only a few tissues, with 93,395 protein pairs having likely associations in all tissues (Fig. 2d).

### Differences between tissues are not primarily driven by gene expression

One of the well-known drivers of differences in protein interactions between tissues is gene expression – the expression of a gene allows for its protein to be present and interact in a tissue. Indeed, the proteins that were quantified in a given tissue were generally enriched for genes with elevated expression for that same tissue, but not the other tissues (Fig. S7). Likewise, the associations between proteins with elevated expression profiles for a tissue were more likely than their associations with all proteins (association scores 0.58 ± 0.05 vs 0.34 ± 0.01 respectively). However, the number of likely associations between proteins with elevated expression profiles remained limited to ∼0.06% of all likely associations of the tissues (Fig. S7), demonstrating that differences in gene expression are reflected in our association atlas but do not define the likely associations for each tissue.

We further explored the variation of likely associations between pairs of tissues. Specifically, for a given tissue, we found that on average, 22.2% of likely associations were shared with other tissues, while 10.8% were unlikely in other tissues and 30% were not quantified in other tissues (Fig. 2e). We then asked to what extent these differences are explained by gene expression. We found that, on average, 50.9% of the unquantified associations involved proteins that were not expressed or whose expression was reduced by at least 2-fold in the other tissue (representing 14.2% of all likely associations). Similarly, 31.8% of associations that are unlikely in other tissues involved proteins with reduced expression (3.4% of all likely associations). Thus, only around 18% of differences in likely associations between tissues are explainable by changes in gene expression. Finally, using the healthy- and tumor-derived scores as independent replicates for the same tissue, we found that, on average, 52.3% of likely associations were reproducible, while 7.3% and 1.1% of likely associations were not reproducible by being unlikely or unquantified respectively. Given that 52% of likely associations reproduce between replicates, and that on average 22% of likely associations are also likely in other tissues, we estimate that up to 30% of likely associations are tissue-specific (Fig. S8).

### Tissues share known interactions and reveal tissue-specific cellular components

Finally, we sought to characterize the likely associations that were shared between tissues. Compared to all likely associations (average Jaccard index ∼0.12), we found that the similarity between pairs of tissues increased as we restricted the associations to interactions identified through high-throughput screens such as yeast two-hybrid (Jaccard index ∼0.19 - HuRI [3]) or affinity purification (∼0.34 - BioPlex [4]) experiments (Fig. 2f). Likewise, the similarity between pairs of tissues increased when restricting likely associations to known interactions reported for signaling (0.24 - SIGNOR [78]), biological pathways (0.37 - Reactome [79]), functional associations (0.46 - STRING [80]) or well-defined human protein complexes (0.67 - CORUM). Our association atlas thus reflects the characteristics of protein interactions, with functional and signaling interactions being less commonly shared between tissues than protein complexes.

Well characterized protein complexes were generally preserved across tissues, becoming more variable as the complex-averaged association scores decreased (rho = −0.77, −log10-p-val = 125; Fig. S8). As seen in other proteomics datasets [81], more variable complexes typically involved signaling and regulation (e.g., TNF and Emerin) while more stable complexes involved central cellular structures (e.g., ribosomes and the respiratory chain – Fig. S9). While protein complexes varied little between tissues, we found that protein associations varied strongly for tissue-specific cellular components (Gene Ontology) of the brain (synapse-related components), throat (structural components of muscle fiber), ovary (e.g., ER luminal membrane), lung (e.g., 9+2 motile cilium) and the liver (e.g. microbodies) (Fig. S10). This suggests that tissue and cell-type specific cellular components are an important driver of tissue-specific protein interactions that are independent of simple expression differences.

### Association atlas reveals cell-type specific associations

To explore cell-type specific associations in our association atlas, we took the AP-2 adaptor complex as a well-known example. The AP-2 complex has neuron-specific functions besides functions that are general to all cells [82]. Indeed, the subunits of the AP-2 complex were co-abundant in all tissues (average association score between subunits ∼0.80). We found 103 proteins that had association scores with all AP-2 subunits in all tissues and were known to associate with AP-2 (STRING score > 400). Among these, the 37 synaptic proteins (SynGO [83]) had higher association scores with the AP-2 complex in the brain (average 0.58) compared to the other tissues (0.5 ± 0.01; p-value = 0.027 – MWU-test). Conversely, the non-synaptic interactors had lower association scores with the AP-2 complex in the brain (0.32) compared to the other tissues (0.46 ± 0.01; p-value 5e-7) (Fig. 3a). We explored further examples by focusing on cell-type specific associations in the context of disease. We found that many proteins of hemoglobin are related to anemia and have likely associations with anemia proteins, but only in the blood (Fig. 3b; Methods). Likewise, we found that subunits of chylomicron - which transports dairy lipids from the intestines – contain and have likely associations with proteins related to Crohn’s disease, but only in the colon [84,85]. Finally, we found that subunits of fibrinogen– synthesized in the liver – contain and have liver-only likely associations with proteins related to liver disease [86,87]. For the other tissues, we could find many examples of cell-type specific associations of protein complexes or cellular components that related to tissue-specific disorders such as diabetes and kidney disease (Fig. S11). These examples demonstrate that our association atlas can be used to study tissue-specific functions of protein complexes and context-dependent associations for disease genes.

**Figure 3.**
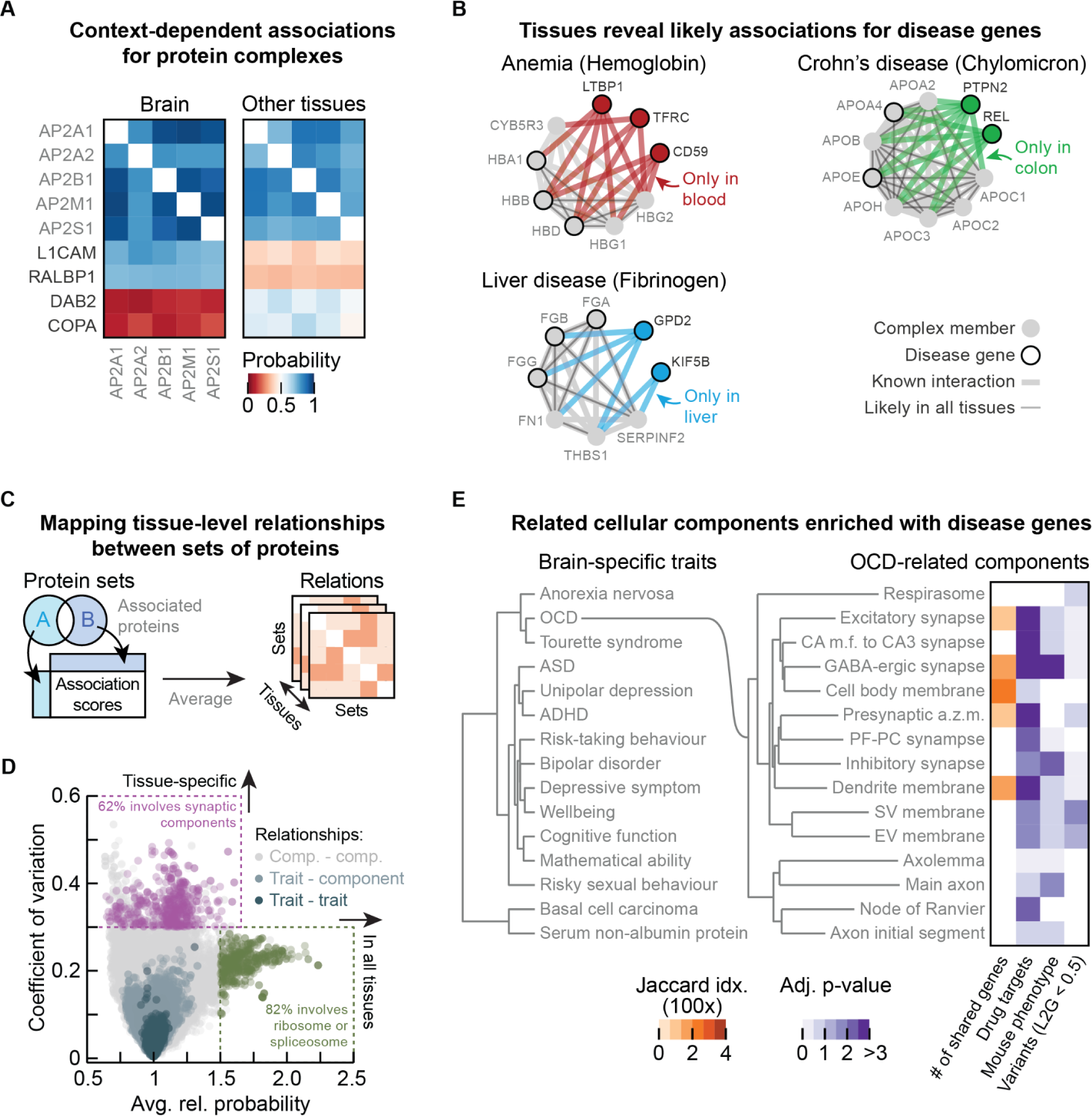
Association scores define relationships between protein sets. **(a)** Association scores between AP-2 subunits and with known AP-2 interactors (STRING scores > 400) that are synaptic proteins (L1CAM, RALBP1 - SynGO) or not (DAB2, COPA). Heatmaps represent the association scores in the brain and the averaged association scores in the other tissues. **(b)** Associations of Hemoglobin (GO:0005833) with proteins associated to anemia, Chylomicron (GO:0042627) with proteins associated to Crohn’s disease, and Fibrinogen (GO:0005577) with proteins associated to liver disease. Proteins (nodes) are complex members (gray) or disease genes (black edge). Associations are likely in all tissues (thin gray lines) or likely in a single tissue and not likely in all others (thick colored lines). Thick gray lines are interactions with prior evidence (STRING scores exceeding 100). Complexes defined through GO annotation, including proteins quantified in all tissues with STRING scores amongst subunits exceeding 750. Disease genes were defined through GWAS variants (OTAR L2G > 0.5). **(c)** Schematic of approach. Proteins associated to pre-defined groups are used to score relationships between sets of proteins by aggregating the association scores of proteins that are not shared between the sets. **(d)** Relationship scores between cellular components (light gray - Gene Ontology), GWAS traits (dark blue - L2G > 0.5) and across traits and components (light blue). Each dot represents the relationship between two protein sets. Shown are the average and coefficient of variation of relationship scores relative to the tissue median. Green dots show relations of the ribosome and spliceosome (for avg. rel. score > 1.5 - green box), purple dots show relations of synaptic components (for CV > 0.35 - purple box). **(e)** Dendrogram (left) of 15 GWAS traits most specific to the brain as measured by z-scores of the median association probability between trait-associated proteins (L2G > 0.5) relative to the tissue median. Dendrogram (right) of 15 GO cellular components having the highest relationship scores with Obsessive-Compulsive Disorder (OCD) in the brain. Heatmaps show overlapping genes between cellular components and OCD (orange - Jaccard Index) and the enrichment (Benjamini-Hochberg (BH) adjusted p-values) of non-overlapping genes from the cellular components with drug targets for OCD (purple - ChEMBL clinical stage II or higher), genes associated to OCD in mice (IMPC), or other GWAS variants associated to OCD (L2G score < 0.5). Dendrograms were constructed with complete-linkage clustering using the Manhattan distance on the relationship scores between traits (left) and the relationship scores between cellular components (right).

### Tissue-specific relationships between traits and cellular components

We sought to generalize these examples of context-relevant associations by systematically mapping the relationships amongst traits and multi-protein structures such as complexes and cellular components. We defined the relationship score between two sets of proteins as the median association score between all pairs of proteins that are not shared between the sets (Fig. 3c – schematic). This methodology thus avoids scoring relationships with the similarity between sets that is simply due to the proteins that they have in common. Protein sets for components were defined based on GO cellular components and protein sets for human traits/diseases were defined based on evidence of their coding genes being associated with traits through GWAS (Open Targets L2G score > 0.5 [88–90]). We computed the relationship scores between traits (37.382 pairs), between traits and cellular components (124.079 pairs), and between components (101.823 pairs; Tables S3-5). Pairs of these protein sets generally shared few proteins (average Jaccard index ∼0.009), with the (non-zero) similarity between protein sets being a poor predictor of relationship scores (Pearson correlation coefficient = −0.22). Relative to the median association scores of the tissues, we found that the relationship scores of cellular components widely varied between pairs of components and across tissues, compared to the relationship scores of the traits (Fig. 3d). Moreover, the relationship scores that were high in all tissues were primarily of core cellular components such as the ribosome and spliceosome (82% of relationships with relative average score > 1.5), while the relationship scores that varied most across tissues often involved tissue-specific compartments such as synaptic components (62% of relationships with CV > 0.35). These observations suggest that the relationship scores recapitulate relatedness of protein sets in a tissue-specific manner, particularly for the brain.

### Relationship scores for prioritizing disease genes

Focusing on the brain, we aggregated the association scores at the trait-level to unbiasedly select the 15 traits whose GWAS-linked proteins had protein-protein associations most specific to the brain (Methods; Tables S6-7). Indeed, 13 of these 15 traits were related to the brain (e.g., neuronal disorders – more general, 58% of the top 50 traits were related to the brain; Table S8). We then clustered these brain-specific traits using the trait-trait relationship scores from the brain, revealing a hierarchical organization of traits with co-occurring conditions such as anorexia nervosa, obsessive-compulsive disorder (OCD) and Tourette syndrome closely clustering together (Fig. 3d – left dendrogram) [91,92]. As an example, using the trait-component relationships, we further determined the 15 cellular components that had the highest relationship scores with obsessive-compulsive disorder (OCD) in the brain, all but two of which were related or specific to neurons (Fig. 3e – right dendrogram). The majority of these cellular components had no genes in common with the genes confidently associated with OCD (Fig. 3d – orange heatmap; Jaccard indices < 0.02). However, after removing the few genes confidently associated with OCD through GWAS, we found that all components were still enriched with or contained OCD-related genes (Fig. 3d, purple heatmaps) – i.e., drug targets for OCD (ChEMBL clinical stage II or higher [93]), genes related to OCD from mice deletion phenotypes (IMPC [94]), or genes less confidently linked to OCD through GWAS (L2G score < 0.5 - Methods). Moreover, these 15 components with the strongest OCD-relationships in the brain were more strongly enriched with OCD-related genes than other cellular components that contained OCD-linked genes (MWU-test p-values: 8.3e-5 (drug targets), 4.9e-4 (genes related to OCD in mice), and 0.03 (genes less confidently linked to OCD through GWAS) - Methods).

Together, the results above demonstrate that the relationship scores can prioritize cellular components that are enriched with trait-relevant genes. Analogously, we found that we could use the relationship scores for reconstructing the hierarchical organization of the cell, including maps of subcellular structures, cellular components, and modules of tissue-specific relations between cellular components (Fig. S12). These observations demonstrate the potential for our association atlas to facilitate the systematic mapping of relations between traits, cellular compartments and likely other ontology terms.

### A network of validated brain-interactions for schizophrenia genes

The results above indicate that the tissue-specific associations could facilitate the prioritization of disease-linked genes by functional association. Indeed, direct interactors of disease linked genes have been used to prioritize causal genes in genetically linked loci and shown to be enriched in successful drug candidates [95–97]. To explore this in more detail we constructed a network of brain-interactions for schizophrenia. We started by taking n=368 genes associated with schizophrenia through GWAS studies (“starting genes” – L2G scores > 0.5) and computed the top 15 traits and cellular components that had the strongest tissue-specific relation to schizophrenia in each tissue (Fig. 4a – Methods). This gave us a collection of genes related to schizophrenia in a tissue-specific manner. We then considered as potential interactions all protein pairs that had one schizophrenia starting gene and one related gene. For each tissue, we filtered these potential interactions for being in the 97-th percentile of the tissue-scores, leading to tissue-specific networks of schizophrenia-associations (Fig. S13; Table S9). After removing the schizophrenia starting genes from these networks, the remaining genes were strongly enriched for genes associated to schizophrenia in mice (log10-p-value ∼17.0 – Fisher’s exact test), drug targets for schizophrenia (∼4.0 – ChEMBL clinical stage II and above), and variants associated to schizophrenia (∼12.3 – OTAR L2G scores < 0.5). This enrichment was specific to the brain compared to any of the other tissues, suggesting that the proposed methodology presents a systematic approach for prioritizing disease genes of tissue-specific traits (Fig. 4b).

**Figure 4.**
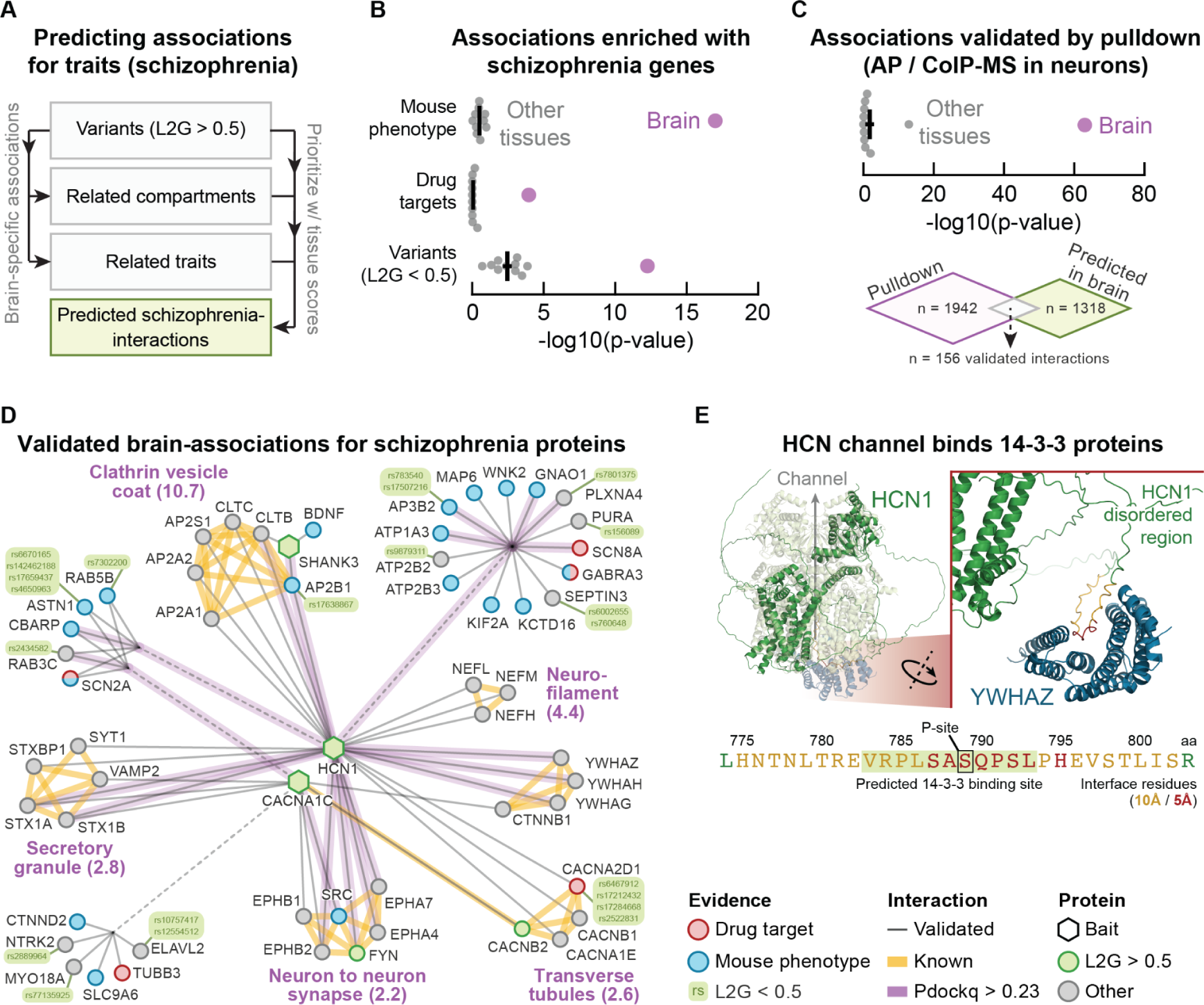
Network of validated brain-associations for schizophrenia proteins. **(a)** Schematic of approach. Genetic variants associated with schizophrenia are used together with relationships between traits and cellular components and the tissue-scores to prioritize potential schizophrenia-interactions. **(b)** Enrichment of predicted brain-interactions with genes associated with schizophrenia through mouse phenotypes (IPMC), drug targets (clinical stage II or higher) or GWAS variants (L2G scores below 0.5). **(c)** Enrichment of pulldown interactions in the predicted brain-interactions of schizophrenia genes. In (b-c), purple dots represent the brain, grey dots are other tissues. Error bars show mean with s.e.m.. **(d)** Simplified network of validated brain-interactions for schizophrenia genes (Methods). Circular and hexagonal nodes were prey and bait proteins in the pulldown studies respectively. Nodes are colored as GWAS variants (green), drug targets (red) or associated with schizophrenia in mice (blue) or other (grey). Grey edges were predicted from tissue scores in the brain and validated through pulldown experiments. Yellow edges are known interactors (i.e., reported in STRING with scores above 750). Purple edges have pdockq scores above 0.23. Purple labels are cellular components most enriched (Benjamini-Hochberg (BH) adjusted p-value between brackets) for genes in the subgraphs of known interactions. **(e)** AlphaFold2 model of the interface between HCN1 and the 14-3-3 proteins (structure of YWHAZ shown). Shown are the HCN complex (cryoEM pdb6UQF - light green) aligned with the AF2 model of HCN (dark green - interface yellow & red) with YWHAZ (blue). Sequence shows the 14-3-3 binding site in HCN1 according to the AF2 models, overlaid with the predicted 14-3-3 binding site (black box - 1433pred score 0.457). Interface residues are colored according to cutoff (10A (yellow), 5A (red)).

Indeed, for brain-related disorders - including autism, ADHD, bipolar disorder, and others - we found that the proposed methodology for predicting disease-related protein associations generally enriched - specifically in the brain - for genes associated to the respective disorder through mouse phenotypes or drug targets, with many also being enriched for GWAS variants that are less confidently linked to the disorder (Fig. S14).

To further validate the networks of predicted schizophrenia-associations, we assembled a curated dataset of brain interactions established experimentally through pulldowns using human brain cells (i.e., AP-MS or CoIP-MS in human micro-dissected brain tissue or human iPSC-derived neurons [98–102]). This dataset contained 7,887 human brain interactions for 30 bait proteins and has been incorporated into the IntAct database [9] (Table S10). We filtered these brain interactions for the bait proteins associated with schizophrenia (OTAR L2G score above 0.5), and further filtered the predicted schizophrenia-associations for ones involving at least one bait protein. The remaining schizophrenia-associations were strongly enriched with interactions for schizophrenia-baits, especially for the brain (log10 p-value ∼63.0 – Fisher exact test) compared to the other tissues (∼1.8; Fig. 4c). Thus, the predicted schizophrenia-associations for the brain - but not the other tissues - are experimentally validated by pulldown studies and enriched with genes related to schizophrenia.

Finally, we combined the predicted schizophrenia-associations for the brain with the pulldown interactions to obtain a network of 156 validated brain-interactions related to schizophrenia, that we simplified to only show genes having prior evidence (Fig. 5d – Methods, Table S11). The visualized network contains 54 proteins connected to 3 bait proteins through a collection of 64 validated brain interactions. These connected proteins include schizophrenia drug targets (5 proteins – clinical stage II and higher), proteins associated with schizophrenia in mice (19 proteins), or proteins linked with schizophrenia with weaker prior evidence (13 proteins – OTAR L2G scores < 0.5). Surprisingly, only two of the validated brain interactions were reported in any of the major protein interaction databases before our curation effort (CACNA1C with CACNB2 and SRC). Moreover, only 18 of the 64 validated brain interactions are quantified in any of the other tissues. Of these 18 interactions, only four have average association scores above 0.5 and only the interactions of SHANK3 with AP2B1 and CLTB are quantified in more than three other tissues (interaction scores of 0.36 ± 0.04 and 0.31 ± 0.02 respectively). These observations support our earlier analysis that the network of validated schizophrenia-interactions is specific to the brain.

**Figure 5.**
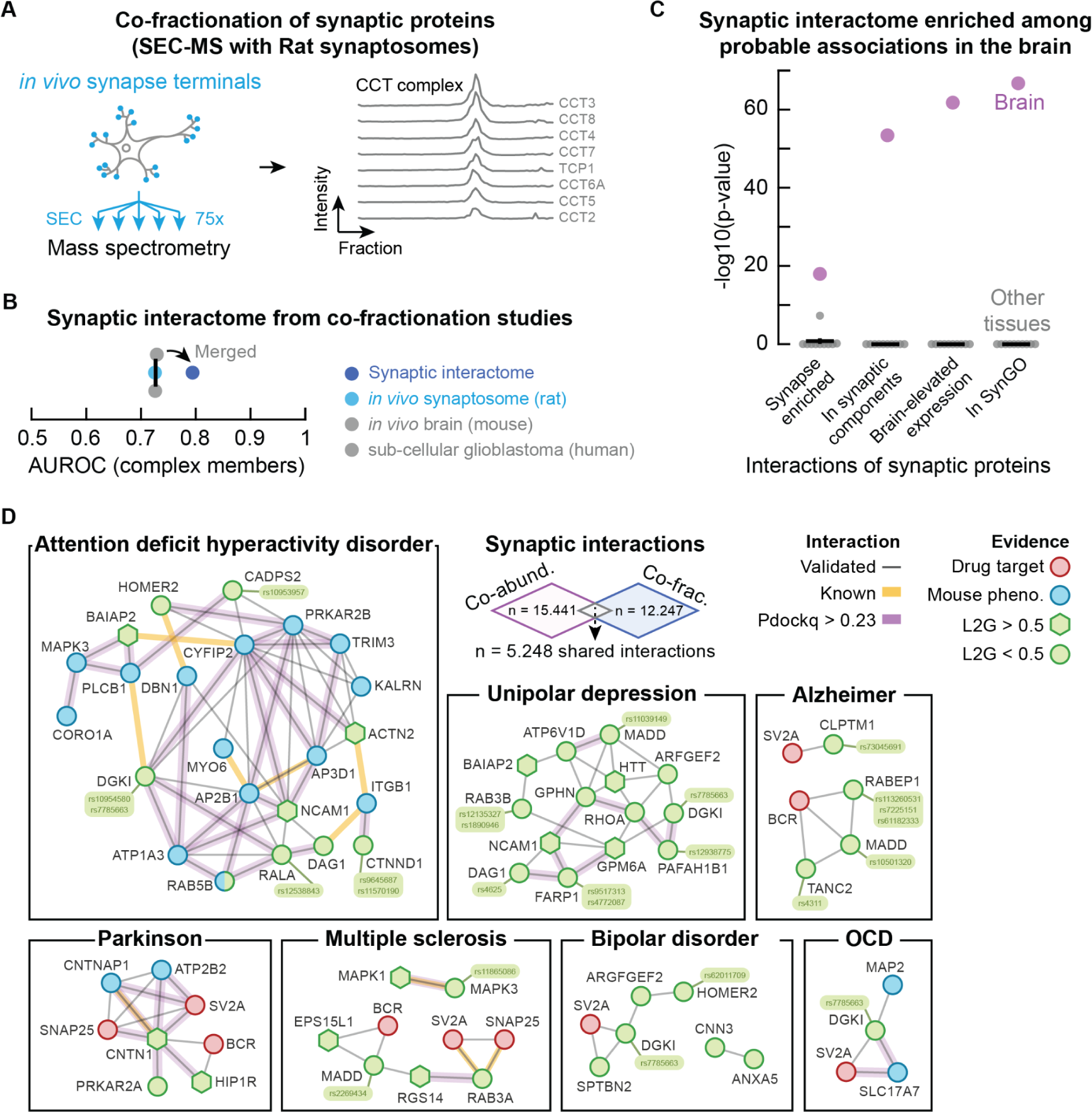
Synaptic interactome. **(a)** Schematic of experiment. *In vivo* synaptosomes (light blue) from rat neurons were purified and fractionated into 75 fractions through SEC and subjected to LC-MS/MS. Elution profiles show the unprocessed protein intensities from mass spectrometry for the CCT complex members (grey). **(b)** AUC values for the co-fractionation studies (*in vivo* mouse brain (gray), subcellular glioblastoma (gray), *in vivo* rat synaptosome (light blue)) and for the merged synaptic interactome (dark blue). Positives defined by complex members in CORUM. **(c)** Comparing the distributions of association scores for interactions of synaptic proteins and for interactions between other proteins from the synaptic interactome. Shown are the p-values (MWU-test) for the brain (purple dots) and other tissues (grey dots). Synaptic proteins: synapse enriched, in SynGO, in GO synaptic components, or brain-elevated expression (at least 2-fold higher expression compared to other tissues on average - Protein Atlas). In (b-c), error bars show mean with s.e.m.. **(d)** Networks of validated synaptic protein associations for genes related to ADHD, unipolar depression, Alzheimer’s disease, Parkinson’s disease, multiple sclerosis, bipolar disorder, and OCD. Shown are associations of synaptic proteins (SynGO) that are co-abundant in the brain and co-fractionated in the synaptosome. Hexagonal nodes are confident GWAS variants (OTAR L2G > 0.5). Circular nodes are colored by association to each disease through drug targets (ChEMBL clinical stage II or higher - red), mouse phenotypes (IMPC - blue) or GWAS variants (L2G < 0.5 - green). Grey edges were predicted from tissue scores in the brain and validated through the co-fractionation studies. Yellow edged are known interactors (STRING scores exceeding 400). Purple edges have an AlphaFold2 model with pdockq-score above 0.23.

The network contains several groups of highly interconnected proteins (STRING score > 750). These groups are enriched with genes of cellular components typical for neuronal functioning and schizophrenia, such as a group of proteins for neurofilaments (log10 p-value 4.4 - [103]) or for clathrin vesicle coat (log10 p-value 10.7). For the clathrin vesicle coat, the network connects all but one subunit of the AP-2 complex and clathrin proteins to HCN1. Interestingly, previous pulldown studies have shown that HCN channels directly interact with TRIP8b [104,105]. TRIP8b regulates trafficking of HCN channels [105] and mainly associates with the AP-2 complex [106]. Moreover, while AP-2 and clathrin are not cell-type specific, both HCN1 and TRIP8b were found to be enriched at PV synapses [107], with HCN channels being specific to PV neurons and important for their high firing frequencies [108,109]. Given the link of PV-neurons with schizophrenia [110–114], these observations suggest that AP-2 and clathrin may be involved in a PV-neuron specific disruption of HCN channel trafficking with schizophrenia.

To suggest putative interface models for the validated schizophrenia-interactions we used AlphaFold2 to predict the structures for 211 protein interactions, including the entire visualized network (Fig. 4d; Table S12). The predicted models were enriched for confident interactions when compared to models for known interactors (e.g., complex members from CORUM [115], p-value = 2.7e-14 - one-sided MWU test). In total, we identified 46 moderate-confidence interactions (pDockQ scores > 0.23). These included the brain-specific binding of SHANK3 to AP2B1 (pDockQ = 0.34; association score ∼0.82 in the brain, compared to ∼0.36 when averaged over the other tissues). Additionally, we found that all three 14-3-3 proteins (YWHAG, YWHAH and YWHAZ) had confident interfaces with HCN1 (Fig. 4e - pDockQ scores exceeding 0.45). The interfaces of these three models overlap and are located in the C-terminal disordered region of HCN1 (residues 775-802). This consensus interface includes a predicted 14-4-3 binding site (centered around S789; 1433Pred score 0.457 [116]) that has been verified through pulldown experiments [117]. Indeed, the binding of 14-3-3 proteins with HCN1 was found to be dependent on the phosphorylation of S789, with the interaction between 14-3-3 and HCN1 likely inhibiting HCN1 degradation [117].

Finally, the network contains 13 genes within loci genetically associated with schizophrenia that had weaker evidence supporting them as the causal genes at each locus. Given their interaction with other schizophrenia associated genes, these could be prioritized as more likely causal due to their functional roles. Some of these genes (AP2B1, ELAVL2 and MYO18A) have the highest locus to gene score for their respective locus with SNPs linked to the schizophrenia but had scores just below the 0.5 cutoff used (0.457, 0.342, 0.428 respectively) [89,90]. In addition to a member of the AP-2 complex (AP2B1) we also found a member of the AP-3 complex (AP3B2). AP3B2 is the second highest linked gene to the locus with the linked SNP (rs783540) having splicing and expression QTL associations with AP3B2 but ranked higher for disruption of CPEB1 given that the variant lies within a CPEB1 intron. Finally, both NTRK2 and PLXNA4 have the second highest L2G score in their locus but the top associated genes (AGTPBP1 and PODXL respectively) have lower or less specific brain expression patterns [12].

### Co-fractionation of synaptosomes to derive a synapse-specific interactome

As a final application of our protein association atlas, we focused on the interactome of synapses. To do so, we prepared and purified synaptosomes from rat brains as an orthogonal approach for validating the brain associations. We fractionated the synaptosomes with size exclusion chromatography (SEC) into 75 fractions that were then subjected to LC-MS/MS. A total of 3,409 unique proteins were detected, including well-known protein complexes such as the CCT complex subunits, whose profiles correlate across the fractions (Fig. 5a - average correlation coefficient of ∼0.96).

We pre-processed the fractionation profiles of the synaptosome and computed the co-abundance of proteins across fractions to score the co-fractionation of 4,200,651 protein pairs in rat synapses (Methods). To increase the confidence of the interaction scores, we then combined the rat synaptosome with other co-fractionation profiles from *in vivo* mouse brains [6] and subcellular fractionation profiles from human glioblastoma cells [7]. We merged the co-fractionation studies by orthologs that were quantified in all three datasets, and computed interaction probabilities with a logistic model using the CORUM database as positives (Methods; Table S13). The resulting synaptic interactome quantified 1,287,210 interaction probabilities for 1,605 proteins, and improved the recovery of known interactions compared to the co-fractionation studies individually (Fig. 5b – AUCs ∼0.79 and ∼0.73 respectively). Of the 1,605 proteins in the interactome, 18% are annotated as synaptic proteins in the SynGO database, 49% have been reported as synapse-enriched in mouse brains, and 56% have previously been identified through cross-linking MS (XL-MS) of the mouse synaptosome [83,107,118]. All XL-MS interactions for these proteins were quantified in our synaptic interactome with ∼64% of these interactions being scored as likely (p-value 1.2e-46 - MWU-test). Moreover, the synaptic interactions were more likely when they involved synaptic proteins compared to the interactions between other proteins (p-value < 6e-159 for SynGO or synapse-enriched proteins - MWU-test). Together, these observations suggest that our synaptic interactome largely aligns with current state-of-the-art synaptic interaction resources and scores the likelihood of interactions for a wide range of synaptic proteins, with interactions being more likely for synaptic proteins compared to other proteins.

### Networks of validated synaptic interactions for brain-disease genes

The interactions involving synaptic proteins enriched for more likely associations compared to non-synaptic proteins, especially for the brain compared to the other tissues of our association atlas (Fig. 5c). Given this brain-specific elevated likelihood of synaptic interactions, we constructed a network of interactions between synaptic proteins (SynGO) that were both likely co-abundant in the brain (15.441 associations) and likely co-fractionated for the synaptosome (12.247 interactions). The resulting network consisted of a collection of 5.248 validated protein interactions between synaptic proteins (Table S14). These synaptic interactions were primarily specific to the brain, since few of the interactions were likely for the majority of other tissues in our association atlas (∼16.8%), and only a small fraction was reported in any protein interaction database (5.7% in STRING>400, 3.7% in HuMAP>0.99 [1], 2.4% in IntAct, and 2% in BioPlex).

We were particularly interested in the validated interactions of synaptic proteins that are associated with brain disorders. As before, we filtered for the 43 GWAS traits that had associated mouse phenotype data (IMPC) or known drug targets (ChEMBL) and whose trait-level association scores were elevated in the brain compared to the other tissues (z-score > 0). We found that 11 of these 43 traits were disorders clearly related to the brain (Table S15). From these 11 traits we selected the 514 synaptic interactions that involve at least one disease gene. Additionally, to suggest putative interface models, we used AlphaFold2 to predict the structures for these 514 interactions with the predicted models being enriched for confident interactions (high pDockQ scores) when compared to models for known interactors (e.g., complex members from CORUM (p-value = 4e-73) or HuMAP (p-value = 3e-5); MWU-test). In total, we identified 201 moderate-confidence interactions (pDockQ scores > 0.23; Table S16). Finally, we visualized the networks of validated synaptic interactions between disease genes for each of the brain disorders (Fig. 5d - Methods).

### Prioritizing synaptic disease genes as more likely to be causal for brain disorders

Several genes in these networks have weaker prior evidence supporting them as the causal genes at loci genetically associated with the brain disorders. These genes could be prioritized as more likely to be causal for the disorders due to their validated synaptic interactions with other genes that are confidently associated with the same disorders. As before, we looked for genes with the highest (below cutoff) locus to gene scores for their respective locus with SNPs linked to the brain disorders. We found likely causal genes for Alzheimer (RABEP1 - score 0.471; CLPTM1 - 0.378; MADD - 0.315), multiple sclerosis (MAPK3 - 0.462; MADD - 0.301), unipolar depression (DGKI - 0.378; MADD - 0.244), attention deficit hyperactivity disorder (DGKI - 0.387; CADPS2 - 0.34; CTNND1 - 0.204), and bipolar disorder and OCD (DGKI - 0.378) [89,90]. Of these genes, all but CTNND1, DGKI and RABEP1 have the highest expression in brain tissue [12].

Additionally, we found genes that did not have the highest locus to gene score for the respective SNP linked to the brain disorders, but still had additional evidence. For example, we found that PAFAH1B1 associates with RHOA in our synaptic interactome, both having weak prior evidence for being causal to depression. PAFAH1B1 plays a role in neural mobility and is required for activation of Rho GTPases such as RHOA [119]. PAFAH1B1 was the second highest scoring gene for the locus with a SNP (rs12938775) linked to unipolar depression. This variant has expression QTL associations with PAFAH1B1 and lies within a PAFAH1B1 intron, despite being more distant to the transcription start site of PAFAH1B1 compared to the highest scoring gene (CLUH). However, PAFAH1B1 is a synaptic protein while CLUH is not, with the expression of PAFAH1B1 being higher and more specific to the brain compared to CLUH [12]. Overall, this example demonstrates how orthogonal approaches targeting sub-cellular structures and individual tissues can provide tissue-specific protein interaction networks and aid in the prioritization of genes likely to be causal for tissue-related disorders.

## DISCUSSION

Despite progress in mapping protein interactions on a large scale, we lack a comprehensive understanding of how these interactions differ across tissues. It has been previously shown that the protein-protein associations derived from correlated protein abundance measurements can be more accurate than those derived from mRNA co-expression [15–18]. Here we show additionally that these associations can be more accurate than those found by correlation analysis of co-fractionation. However, in-line with other reports, we observe that the combination of different co-fractionation datasets can improve the predictions [120] and it’s likely that more advanced analyses of co-fractionation data may yield more accurate results [121]. Interestingly, we note that we can exclude the variation in protein levels that is explained by mRNA changes without significantly affecting the performance of the method. This suggests that protein interactions partners are often post-transcriptionally co-regulated. This is in-line with the observations that protein abundances of subunits within a complex can be limited by the abundance of the complex itself with active degradation of excess unbound subunits [15,16,22,24,122].

Several lines of evidence attest to the quality of the derived association scores: cohorts of the same tissue have similar association scores; association scores derived from tumor and healthy controls from the same tissue tend to replicate in a tissue-specific manner; and essential protein complexes are generally very well predicted across all tissues. Nevertheless, the replication between healthy and tumor of the likely associations is on average around 50%, suggesting that these associations can be useful in generating testable hypotheses regarding tissue-specific biology, for example for measuring associations between proteins assigned to specific compartments in a tissue-specific manner. However, these associations should be complemented with other data, in particular when deriving hypotheses about any given interaction. For the brain network, we have complemented the co-abundance association with experimental data from pull-downs, co-fractionation and AlphaFold2 predictions to provide a more confident subset of interactions for disease gene mapping. Our associations are derived from human tissues and therefore can provide supporting evidence for interactions obtained from less physiologically relevant material.

We estimate that up to 30% of associations are tissue specific. While we observe a large fraction of associations that are not quantified in different tissues, likely due to technical issues of detectability, we estimate that only a small fraction of the differences across tissues are due to changes in expression levels. We find that cell-type specific structures such as synaptic components may contribute significantly to differences in protein associations across tissues, with differences in post-translational modifications being other potential mechanisms that could explain some variation in tissue-specific protein associations.

Finally, we show that our atlas can be used to derive relations between traits and with cellular components. This approach is particularly effective for disorders of the brain and synaptic components, demonstrating that the network of brain associations is more specific and distinct from the other tissues. We present validated brain-interactions enriched for disease associated genes and drug targets, and show how this can be useful to prioritize novel disease linked genes and presumably also for prioritizing drug candidates. The combination of genetics, tissue-specific networks and AlphaFold2 models provide an integrated network for enhanced understanding of disease mechanisms and target prioritization. In addition, our approach may improve safety of targeting disease genes as the predicted disease-gene associations will tend to be tissue specific and therefore the proposed genes may be safer to target.

## METHODS

### Homogenizing protein identifiers across cohorts

Protein abundance measurements were primarily reported at the gene-level and using gene symbols as gene identifiers (45 of 50 cohorts). Gene identifiers were identified, checked, and corrected where possible to maximize the number of overlapping genes between cohorts. Identifiers that contained multiple gene names were split into individual replicate measurements (for 9 of 50 cohorts). All gene identifiers that were aliases, previous symbols, Ensembl IDs or UniProt IDs were converted to the approved HGNC gene symbols (using HGNC dataset and Ensembl Biomart; Table S19). Thus, gene identifiers were not changed if they were already an approved HGNC gene symbol. Identifiers that mapped to multiple approved gene symbols were ignored, and symbols were corrected for common errors made with gene symbols (e.g., replacing ‘Sep.01’ with ‘SEPTIN1’). Finally, identifiers that could not be mapped to approved HGNC gene symbols and that occurred in at least 5 cohorts were kept with their original identifier. Overall, we mapped ∼99.5% of the reported gene identifiers in the cohorts to 16.881 unique gene symbols (Table S20).

### Pre-processing abundance data

Publicly available protein abundance data was filtered by removing non-abundance samples and measurements (e.g., internal controls, reference samples, samples that were withdrawn or that did not have further annotation, etc.). Protein measurements without gene identifier were removed, and the abundance data of cohorts with replicate samples for some but not all patients were averaged at the patient level (for 5 of 50 cohorts, e.g. for cohorts having a sample with 5 technical replicates as internal control). Zeros were replaced with NaNs for all entries of cohorts that were not log-transformed to ensure that undetected proteins did not have abundance data. Cohorts were then split into tumor and healthy samples if not reported separately. The sample names were homogenized to obtain paired RNA expression and protein abundance data and paired tumor and healthy biopsies. The gene identifiers were homogenized across cohorts to allow for comparison between studies (following ‘Homogenizing protein identifiers across cohorts’). Then, sequentially, duplicate measurements were averaged (i.e., averaging the reported abundance of proteins whose identifier occurred more than once), proteins with measurements in fewer than 10 patients were removed, and the data was log2-transformed if not already on a log-scale (13 of 50 cohorts). Finally, the abundance of each protein was median normalized across patients. Note that only the log-transformation affects the co-abundance estimates, since correlation coefficients are invariant under linear transformations. RNA expression data was processed in analogous fashion.

As deviations from the above, samples were removed from cohorts that shared patients by filtering the follow-up studies for patients whose protein abundance was not previously reported [47,48]. Moreover, zeros were replaced with 10% of the smallest non-zero abundance value before performing a log-transformation for cohorts whose data was reported as a fold-change and whose zeros did not represent unmeasured proteins (reported as NaNs; [37]). If applicable (5 of the 50 cohorts), iBAQ intensities were converted to fraction of total values (FOT) by dividing the iBAQ intensities by the total iBAQ (sum over samples and proteins) and multiplying all values by 10e5, after which proteins were removed when the FOT did not exceed 10e-5 for any of the samples.

### Computing association probabilities

Co-abundance estimates were computed by calculating the correlation matrix consisting of Pearson correlations between all possible pairs of proteins for a cohort. Each pair of proteins was required to have at least n = 30 paired abundances across samples for computing a correlation value (i.e., protein pairs must be quantified in at least 30 of the same samples - available for 48 out of 50 studies). Correlations were then reported for each unique protein pair (alphabetically sorted by gene symbols), ignoring self-correlations and missing values. Finally, correlations were converted to association probabilities as follows. Protein pairs were annotated as being complex members (co-occuring in a protein complex in the CORUM database [76]) or not. A logistic regression was then fitted to the correlation values using the complex members as positives and all other protein pairs as negatives. Positive and negative classes were balanced by adjusting the weights of protein pairs inversely proportional to the class frequencies (balanced class weights). The model included an intercept and no penalty for parameter values was used. Finally, the fitted logistic model was used to transform the correlation values to probabilities. Note that the model only transforms correlations (in the range −1 to 1) to probabilities (range 0 to 1) and thereby yields the same ranking of protein pairs when sorting correlations and probabilities from large to small. The association probabilities per tissue are available through the BioStudies repository for this paper https://www.ebi.ac.uk/biostudies/studies/S-BSST1423.

### Computing interaction probabilities from co-fractionation data

Co-fractionation data was collected from studies of in vivo mouse tissues (SEC-PCP-SILAM – 2×55 fractions for 7 tissues) [6], human breast tissue-derived cell lines (IEX-HPLC-MS – 3×2×192 fractions of 3 biological replicates with each 2 technical replicates) [77], and subcellular fractionation of human cancer cell lines (HiRIEF-LC-MS - 5×10 fractions for 5 different tissues, and additionally 4×10 fractions from 3 replicates and 1 condition in the lung) [7]. The protein identifiers of the mouse data were converted to human orthologs (Ensembl BioMart, date = 2023-09-28). Replicates and conditions were merged together and treated as different fractions. The resulting 10 co-fractionation datasets (7 mouse tissues, 1 human breast cell line, 1 human lung cell line and 1 human mixed cell lines) were further processed as follows. Data was filtered for fractions and proteins that had non-zero measurements, and abundances of duplicate protein identifiers were averaged. Empty fractions of each fractionation-experiment were then set to 90% of the minimum non-zero value for each protein, log2-transformed (mouse tissues and breast cell line) and median normalized across samples. Setting empty fractions to 90% of the minimum non-zero value was advantageous as it increased the number of protein pairs having sufficient observations across fractions for computing association probabilities (i.e., on average ∼7.5-fold more protein pairs), and improved the co-fractionation estimates by including the fractions where proteins are co-absent (increased AUCs on average by ∼9.5%). Here, the co-fractionation estimates were not sensitive to the value used for empty fractions (i.e., on average ∼0.5% difference in AUC values between using 90% or 10% of the minimum non-zero values). Interaction probabilities were then computed following ‘Computing association probabilities’ using the correlation of intensities of protein pairs across fractions.

### Computing AUC values

Positives were defined as interactors predicted through prior evidence (e.g., complex members reported in CORUM). Sets of non-duplicate negatives of the same set size as the positives were sampled from all other protein pairs having association scores. AUC values were then computed as the average of the AUC values for the (n = 10) replicate sets of negatives. For each cohort, the same sets of negatives were used to compare the matched association scores from gene expression and protein abundance data.

### Scoring recovery of known complex members from RNA expression, protein abundance, and protein co-fractionation data (Fig. 1d-e)

Studies were selected for having both RNA expression and protein abundance data available (32 studies - 29 studies had sufficient shared samples). For each of these studies, both the RNA and protein abundance data was filtered for genes and samples quantified for both modalities. The RNA expression was then regressed out of the protein abundance data to exclude the variation of protein abundance that can be explained by gene expression (‘protein - RNA’). Specifically, a linear model (linear regression with intercept) was fitted for each protein to find the dependence of protein abundance on RNA expression, ignoring missing values. The gene expression was then regressed out of the protein abundance by subtracting the model’s prediction from the protein abundance for all values. The Pearson correlation coefficient between the RNA expression and protein abundance was zero after regressing out the RNA expression. Co-abundance estimates and association probabilities were computed following ‘Computing association probabilities’. Additionally, for the same studies, the RNA expression and protein abundance data were combined to predict association probabilities (‘protein + RNA‘). Specifically, the correlation estimates for both the RNA co-expression and protein co-abundance of the studies were filtered for protein pairs quantified for both modalities. The correlation estimates were then converted to association probabilities using a logistic model and the two modalities as variates. Complex members from CORUM were used as positives (following ‘Computing association probabilities’).

For each study, the protein pairs were filtered for having association probabilities for both modalities and their combinations (from ‘Combinations of gene expression and protein co-abundance data’). AUC values were then computed following ‘Computing AUC values’ for the filtered sets of association probabilities, using complex members reported in CORUM as positives. AUC values were also computed for the co-fractionation studies (from ‘Computing interaction probabilities from co-fractionation data)’. Protein pairs of co-fractionation studies were not filtered as they were separate studies from the RNA co-expression and protein co-abundance studies.

### Recovering tissue-specific associations (Figs. 1g & 2c)

Cohorts were filtered for protein pairs having association probabilities in all cohorts. The recovery of tissue-specific associations was then scored through a hold-one-out methodology as follows. Each cohort was sequentially used as a hold-out by not using its association scores for predicting tissue-specific associations. Using all other cohorts, tissue-level association probabilities were computed, for each tissue, as the averaged probabilities of the cohorts. Associations of protein pairs were defined as tissue-specific if the tissue-level association probability exceeded the 95-th percentile for a given tissue and remained below 0.5 when averaged over all other tissues. The with-held study was then used to recover the tissue-specific associations of each tissue following ‘Computing AUC values’, resulting in AUC values for all pairs of studies and tissues (Table S21). Recovery of tissue-specific associations for a tissue with the cohorts from some (other) tissue was then summarized at the tissue-level by averaging the AUC scores of the respective cohorts.

Tissue-specific associations were recovered for the tissue-level scores from tumor- and healthy-derived biopsies (Fig. 2c) in analogous fashion. Tissues were filtered for protein pairs having association scores across all studies for both the tumor- and healthy-derived scores. Tissue-specific associations were predicted using the association scores derived from healthy samples. Here, the tissue-specific associations were defined as the protein pairs whose association scores exceeded the 95-th percentile in one tissue and remained below 0.5 when averaged over the other tissues. For each tissue, these tissue-specific associations were then recovered using the association scores derived from the tumor samples.

### Defining protein association atlas for human tissues (Fig. 2a-d)

Association scores for a tissue were computed by averaging, for each protein pair, the association probabilities of all cohorts of the tissue. Missing values were ignored. Three cohorts were excluded based on their performance for recovering tissue-specific associations of the other cohorts of their respective tissues (Fig. S4). Tissues were only included if association probabilities from at least two studies were available (not for the bladder, prostate, uterus (tumor), and only for the lung, throat, liver, kidney, colon and stomach (healthy)). Likely associations were defined as the protein pairs whose association score exceeded 0.5 for a given tissue, confident associations were defined as the protein pairs whose association score exceeded the 99-th percentile for a given tissue.

### Differences of likely associations between tissues (Fig. 2e)

For each pair of tissues, the likely associations of one tissue were classified as being likely, unlikely or not quantified in the other tissue. Here, unlikely associations were protein pairs whose association score was below 0.25 for a tissue. A likely association was classified as not quantified if the protein pair did not have an association score in the other tissue. The likely associations of each tissue classified in all other tissues were then expressed as a percentage of the total number of likely associations of that tissue. Finally, the percentage of associations in each class was averaged over all pairs of tissues. Associations were analogously classified for the six pairs of tumor- and healthy-derived scores.

The likely associations of each tissue that were unlikely or not quantified in other tissues were further classified by gene expression. First, gene expression levels across tissues were obtained from the protein atlas (consensus normalized expression) [12], and aggregated at the whole-tissue-level by taking the maximum expression for each protein over the sub-tissues for the brain (cerebellum, amygdala, cerebral cortex, basal ganglia, choroid plexus, hippocampal formation, hypothalamus, midbrain, pituitary gland, spinal cord) and colon (colon, duodenum, small intestine). Expression of bone marrow was used for blood. For a pair of tissues, the proteins were selected whose expression was at least 2-fold higher in one tissue compared to the other tissue. The likely associations of that tissue that were unlikely or not quantified in the other tissue and that involved at least one of the proteins with 2-fold higher expression were then labeled as being unlikely or not quantified with reduced expression. Analogously, likely associations of one tissue that were not quantified in another tissue were labeled as not quantified without expression if the protein was not expressed in the other tissue (expression level is 0 in the other tissue - these were not considered for the proteins having at least 2-fold lower expression).

### Similarity of likely associations between tissues (Fig. 2f)

For each tissue, the likely associations were filtered for having prior evidence. Specifically, associations were filtered for interactions reported in HuRI (date = 23-02-27) [3], BioPlex (combined interaction networks for 239T, HCT116 (both 22-09-22) and RPE1 (23-04-05)) [4], Signor (23-04-05) [78], Reactome (23-04-05, [79]), STRING (with scores >= 400 – 22-10-24) [80], and all pairwise complex members from HuMAP with scores exceeding the 99-th percentile (23-11-20) [1] and CORUM (22-09-20) [76]. The similarity of likely associations between each pair of tissues was then computed as the Jaccard index of their sets of likely associations after filtering for the different types of prior evidence, or without any filtering.

### Context-dependent associations for protein complexes (Fig. 3a)

Association scores were filtered for protein pairs quantified for all tissues. Proteins were filtered for having associations with all subunits of the AP-2 complex, and further filtered for the 103 proteins that were known to interact with at least one AP-2 subunit (STRING scores > 400). These 103 known interactors were then divided into the 37 synaptic and 66 other (non-synaptic) proteins (following SynGO, release 2021-02-25), and their association scores with the AP-2 subunits were averaged at the complex level (Table S22). The distribution of complex-level association scores for the synaptic proteins were then compared between the brain and the other tissues on average (one-sided MWU-test). The distributions for the non-synaptic proteins were compared following the same methodology. The AP-2 complex was visualized with the interactors that had the largest complex-level difference in association scores between the brain and the other tissues, taking synaptic interactors that had higher association scores in the brain and non-synaptic interactors that had higher association scores in the other tissues.

### Proteins associated with GO cellular components

Descriptions of GO terms were downloaded from Gene Ontology (http://purl.obolibrary.org/obo/go/go-basic.obo - date = 23-06-28 [88]) and filtered for not being obsolete, having a description, and being cellular components but not protein complexes (namespace = ‘cellular_component’, name does not contain ‘complex’ – 2.059 compartments). The cellular compartments were annotated with associated genes (HGCN gene symbols) using the QuickGO annotations from EBI (date = 23-07-03). Additionally, the cellular compartments were annotated with associated genes reported by UniprotKB (date = 23-06-28) through their Rest API. For both sets of associated genes, the cellular compartments were filtered for having at least 10 and at most 500 genes quantified through the association atlas, and were then merged GO-term level. The resulting 454 GO cellular compartments each have at least 10 associated genes from either EBI or UniprotKB reported in the association atlas (Table S23).

### Proteins associated with human traits through GWAS variants, drug targets or mouse phenotypes

Evidence was downloaded from the Open Targets repository (OTAR, date = 23-07-10) [89,90]. Three types of evidence collected from OTAR were used for associating genes to traits. Specifically, genes were considered associated with traits through GWAS, mouse phenotypes (International Mouse Phenotyping Consortium (IMPC)) [94] or as drug targets (ChEMBL) [93]. For genes associated with traits through GWAS, the evidence was loaded from the OTAR ‘ot_genetics_portal’ source (scores exceeding 0.5 were considered as confidently associated genes). Gene identifiers were converted to HGCN gene symbols following the symbols used in the association atlas (following the same identifiers used in ‘Homogenizing protein identifiers across cohorts’). Trait identifiers (EFO) were converted to trait names using annotations from the Open Targets platform. For genes associated with traits through mouse phenotypes and drug targets, the evidence was loaded from the OTAR sources ‘impc’ (scores exceeding 0.5) and ‘chembl’ (clinical stage I or higher) respectively. All gene sets were filtered for genes that were quantified in the association atlas (Tables S24-26).

### Disease-relevant associations for protein complexes (Fig. 3b)

The association scores were filtered for protein pairs quantified for all tissues. For associations with disease genes (Fig. 3b), the disease genes for Anemia (blood), Crohn’s disease (colon) and Liver disease (liver) were then selected by merging together the genes associated to each disease through GWAS (OTAR L2g > 0.5), drug targets (ChEBML clinical stage I or higher) or mouse phenotypes (IMPC). Protein complexes for each disease (Hemoglobin (GO:0005833 - for blood), Chylomicron (GO:0042627 - for colon) and Fibrinogen (GO:0005577 - for liver)) were filtered for complex members that had interactions with the other subunits according to STRING (scores > 750). Tissue-specific associations were then selected for each tissue-related pair of complex and disease, defined as the associations that were likely in the respective tissue but whose score remained below 0.5 in all other tissues. Exemplary disease genes were selected by taking the proteins that had at least two (liver) or four (blood, colon) tissue-specific associations and whose specificity was largest on the complex-level. Specificity of the tissue-specific associations was computed as the difference between the highest (respective tissue) and second highest (other tissues) association score, averaged over the subunits of the protein complex.

### Trait- and component-level association scores and relationship scores of traits and components

Protein sets were defined by the genes associated to GO cellular components (454 components - ‘Proteins associated to GO cellular components’) or genes associated to traits through GWAS (274 traits - ‘Proteins associated with human traits through GWAS variants, drug targets or mouse phenotypes’; traits having at least 50 associated proteins). Set-level association scores were then defined as the median of the association scores between all pairs of proteins in the set that were quantified for a given tissue. Set-level association scores were computed for all protein sets in all tissues (Tables S6-7). The relationship score between protein sets A and B for a given tissue was defined as the median of all association scores between proteins from A (that were not in B) and proteins from B (that were not in A). Relationship scores were then computed for all pairs of protein sets in all tissues (Tables S3-5).

### Specificity and preservation of relationship scores (Fig. 3d)

For each tissue, the relationship scores between cellular components (from ‘Trait- and component-level association scores and relationship scores of traits and components’) were normalized with the median association score of the tissue, and filtered for relationships whose score was quantified for all tissues. Relationships were then filtered for pairs of components whose score averaged across tissues exceeded 1.5, and for which the name of at least one component contained ‘ribosom’, ‘spliceo’ or ‘snRNP’. Alternatively, relationships were filtered for pairs of components whose coefficient of variation across tissues exceeded 0.3 and for which the name of at least one component contained ‘synap’, ‘axo’, ‘neuro’, ‘dendrit’, ‘node of Ranvier’ or ‘calyx of Held’.

### Relationship scores for prioritizing genes associated with OCD (Fig. 3e)

The trait-level association scores (from ‘Trait- and component-level association scores and relationship scores of traits and components’) were normalized with the tissue-median for each tissue and then z-scored across tissues. The top 15 traits most specific to the brain were selected by taking the traits with the highest z-score for the brain tissue (Table S8). The trait-trait relationship scores of the brain were then filtered for all relationships of the brain-specific traits. The resulting matrix scored the relationships of the brain-specific traits with all other traits, and was used to cluster the brain-specific traits. Next, the cellular components that had relationship scores were filtered for components having at least 15 proteins quantified in the association atlas (384 components). The trait-component relationship scores of the brain were filtered for all relationships between these components and OCD, from which the 15 components with the highest relationship scores with OCD were selected (Table S27). The component-component relationship scores of the brain were then filtered for all relationships of these OCD-related components and used for clustering the OCD-related components.

Enrichment of OCD-associated genes in the OCD-related components was then tested with one-sided Fisher’s exact tests. The genes confidently associated with OCD through GWAS (OTAR L2G > 0.5) were removed from all gene sets before testing (these genes were already used to score the relationships between OCD and cellular compartments). For each OCD-related component, the genes in all other cellular components were used as backgrounds. OCD-associated genes were defined as OCD drug targets (ChEMBL clinical stage II and higher), genes associated with OCD in mice (IMPC) or genes with weak evidence supporting them as causal for OCD (OTAR L2G < 0.5 - disorders with at least 10 associated proteins). The collection of all other genes reported in each database was used as a background respectively. Finally, the enrichment of OCD-associated genes in all other cellular components was tested in similar fashion (Table S28). For each database of OCD-associated genes, the cellular components were filtered for the ones that contained OCD-associated genes. The level of enrichment of OCD-associated genes in OCD-related components was then compared with the enrichment in the other components by comparing the distributions of non-adjusted p-values (one-sided MWU-test).

### Predicting associations of schizophrenia genes (Fig. 4a)

For each tissue, a tissue-specific distance matrix of GWAS traits was created by computing the pairwise Manhattan distance between pairs of traits using the relationship scores between traits after z-scoring across tissues. The schizophrenia-related traits were then selected as the top 15 traits closest to schizophrenia. Analogously, the schizophrenia-related GO cellular compartments were selected by z-scoring the relationships between schizophrenia and components across tissues, and then selecting the top 15 components most specific for schizophrenia in each tissue. The potential interactions were then the protein pairs that had one gene confidently associated with schizophrenia through GWAS (L2G scores > 0.5 - n = 368) and one gene from the tissue-specific schizophrenia-related traits and compartments. These potential interactions were filtered for associations whose likelihood exceeded the 97-th percentile of association scores (Table S9).

### Schizophrenia-associations enriched with schizophrenia genes (Fig. 4b)

The 368 genes confidently associated with schizophrenia through GWAS were removed from the collection of proteins involved in the potential schizophrenia-interactions for each tissue. Enrichment of other schizophrenia-associated genes was then tested with a one sided Fisher’s exact test using all other proteins of the associations for each tissue as background. For these tests, the schizophrenia-associated genes were defined as schizophrenia drug targets (ChEMBL clinical stage II and higher), genes associated with schizophrenia in mice (IMPC) or genes with weak evidence supporting them as causal for schizophrenia (OTAR GWAS L2G < 0.5). The collection of all other genes reported in each database were used as background respectively.

### Schizophrenia-associations validated by pulldown interactions (Fig. 4c)

Brain interactions were collected and pooled from pulldown studies using micro-dissected human brain tissue or human iPSC-derived neurons (Table S10) [98–102]. These pulldown interactions were filtered for bait proteins that were confidently associated with schizophrenia (OTAR L2G score > 0.5), and the predicted schizophrenia associations were filtered for associations that involved the same bait proteins. This ensured that the predicted schizophrenia associations were the associations that could have been pulled down with the available baits, and that the pulldown interactions represented the collections of interactions that would have been found for baits available in the association atlas. Enrichment was then tested with a one sided Fisher’s exact test, comparing the predicted schizophrenia associations for the bait proteins with all other associations as background for each tissue respectively. Background for the pulldown interactions were all associations of each tissue that could have been pulled down for the same baits.

### Network of validated interactions for schizophrenia genes (Fig. 4d)

Potential schizophrenia-associations from the brain tissue (n = 1.318) were filtered for interactions reported in the pulldown studies (n = 1.942). The remaining n = 156 protein interactions were defined as the validated interactions for schizophrenia genes. The resulting network of validated interactions was further annotated with the association scores for the other tissues in the association atlas, and whether the interactions were reported in the major protein interaction databases (STRING score > 400, HuMAP interactions exceeding the 99-th percentile, and interactions from HuRI, BioPlex, and IntAct [9]) (Table S11).

### Visualizing the network of validated schizophrenia interactions (Fig. 4d)

The STRING network was filtered for fully connected subgraphs (cliques) of proteins in the network of validated schizophrenia interactions (schizophrenia network). Cliques were selected if they consisted of at least 3 proteins (and all STRING scores exceeded 950) or if they consisted of at least 4 proteins (all STRING scores exceeded 750). The selected cliques were the starting graph of the visualized graph. To this graph, all interactions between the cliques and bait proteins in the schizophrenia network were added, together with any interaction between these bait proteins and other proteins in the schizophrenia network that had prior evidence for being related to schizophrenia (i.e., schizophrenia drug targets (ChEMBL clinical stage II or higher), associated with schizophrenia in mice (IMPC), or had weaker evidence supporting them as causal for schizophrenia (OTAR L2G < 0.5)) (Table S11). Finally, each GO cellular component was tested for being enriched with the genes of each clique (excluding bait) in the visualized network using a one-sided Fisher’s exact test. All other proteins of the brain associations and all other genes reported for the GO cellular components were used as backgrounds. Benjamini-Hochberg adjusted p-values were log-transformed and sorted for the cellular component with the strongest enrichment. Cellular components are shown for adjusted p-values < 0.01.

### Structural models of protein-protein interaction interfaces (Fig. 4-5)

Structures were predicted using AlphaFold-multimer v. 2.3.1. Input protein sequences were downloaded from Uniprot (release 2023_03 to release 2024_02). The use of templates was turned on. Five structures were predicted for each protein pair, and the predicted structure with the highest model confidence score was considered the “best” structure for each pair. The predicted DockQ (pDockQ) scores were calculated for the top ranked structure using the implementation from [115]. Interaction interfaces were defined by considering that any residue with at least one atom within 10 Å (and 5 Å (Fig. 4e)) of the other protein chain is part of the interface.

### Synaptosome preparation

Cortices of adult rats (Sprague–Dawley) were dissected and a synaptosome preparation was generated using Syn-PER (Thermo Scientific) according to the manufacturer’s instructions. All animal experiments were carried out under institutional guidelines (ZH172/18 Kanton Zürich Gesundheitsdirektion Veterinäramt).

### Cell lysis

The procedures used for cell lysis and SEC-based fractionation followed experimental protocols previously published [123,124], with slight adaptations regarding the lysis, detergent and used SEC-column. In short, synaptosomes were lysed in HNN buffer (50 mM HEPES, 100 mM NaCl, 50 mM NaF) supplemented with Protease inhibitor (1x out of a 500x stock solution), 20 mM Sodium-Vanadate (of a 200 mM stock solution), 1 mM PMSF (from a 100 mM stock solution) and 1% n-Dodecyl β-D-maltoside (DDM), from a freshly prepared 10% DDM stock solution. Samples were lysed by gentle pipetting up- and down. The lysate was pre-clarified by centrifugation at 16,000 g at 4 °C for 20 minutes. The supernatant was split to multiple ultracentrifuge tubes in order to remove insoluble cell debris by ultracentrifugation in a TLA 100.1 rotor (Beckman-Coulter) at 35,000 g at 4°C for 15 minutes. Next, supernatants were transferred to Eppendorf tubes and protein concentration was assessed by BCA assay (ThermoFisher Scientific). A total protein amount of 1.2 mg (or 319 μL of protein lysate) was then diluted up to 4 mL with HNN buffer (without detergent). To concentrate the sample and remove surplus detergent, samples were concentrated over a 30 kDa molecular weight cutoff filter (Amicon). Samples were centrifuged at 3,600 g at 4 °C for 5 to 10 minutes, with 5 minutes intervals to monitor lysate volume. First samples were concentrated to approximately 500 μL before diluting them again up to 1 mL. This dilution step was repeated for a total three times, before samples were concentrated to approximately 100 μL.

### Separation of complexes by size-exclusion chromatography (SEC)

For the separation of complexes, a 1260 Infinity II HPLC system (Agilent) connected to a SRT-C SEC-1000 (Sepax) 7.8×300 mm column with a SRT-C SEC-1000 (Sepax) 7.8×50 mm pre-column was employed. The system was operated at 500 μL/min flow rate. The column was equilibrated with SEC-buffer (50 mM HEPES, 100 mM NaCl, pH 7.5) for at least 4 column volumes (CV). Before collection of the separated sample, a replicate of the rat synaptosome extract was injected to prime the column. The input for the prime and SEC-experiment was kept at 950 μg total protein amount (according to BCA). The sample was fractionated into 75 x 100 μL fractionations. For separation the HPLC was cooled to 4°C and the column was placed on ice. Column performance was monitored by injecting before and after the experiment a standard protein mix (Phenome protein standard mixture, diluted 1:3 in SEC-buffer).

### Preparation of SEC-fractions by FASP protocol

SEC-fractions were further processed employing a high-recovery and high-throughput FASP protocol (Fossati et al., Methods Mol Biol, 2021). In short, the 96-well Acroprep advance filter plate (10,000 MWCO) was equilibrated by flushing the plate twice with 100 μL H_2_O (HPLC-grade) followed by centrifugation at 1,800 g. Each fraction was diluted 1:1 (v/v) with SEC-buffer and added to the filter plate. Additionally an aliquot of SEC-input was added to the filter plate. Sample was loaded by centrifugation at 1,800 g to completely remove the SEC-buffer. Next, 50 μL of TUA buffer (5 mM TCEP, 8 M Urea, 20 mM Ambic) was added to each fraction. The plate was incubated for 30 minutes at 400 rpm at 37 °C. After cooling down the plate to room temperature, 20 μL of 35 mM IAA was added to each well and the plate was incubated for 1 hour at 400 rpm at room temperature. Urea, IAA and TCEP were removed by centrifugation at 1,800 g and the plate was washed 3 x 100 μL of 20 mM Ambic with centrifugation steps in-between. After the last centrifugation there should be less than 10 μL 20 mM Ambic left in each well. Before adding the digestion mix, the filter plate was moved to a new receiver plate. Proteins were digested overnight at 300 rpm at 37 °C by adding to each well 1 μg Trypsin and 0.3 μg Lys-C diluted in 50 μL 20 mM Ambic. Peptides were collected by centrifugation at 2,400 g for 30 minutes. The filter plate was subsequently washed with 100 μL of HPLC-grade H2O. To increase recovery of hydrophobic peptides, 50 μL of 50% Acetonitrile was added to the receiver plate. Each fraction was then transferred to LoBind tubes and peptides were dried under reduced pressure at 42 °C.

### MS-data analysis of SEC-fractions

For MS analysis, peptides were reconstituted in 5% acetonitrile and 0.1% formic acid containing iRT peptides (Biognosys). The peptides were analyzed on a Fusion Lumos mass spectrometer (Thermo Scientific) using a 2 cm Acclaim™ PepMap™ 100 C18 HPLC trap column (Thermo Scientific) and 25 cm EASY-Spray™ HPLC analytical column (Thermo Scientific) set-up connected to an EASY-nLC 1200 instrument (Thermo Scientific) as previously described [125,126]. Peptides were loaded in 100% buffer A (98% H2O, 2% acetonitrile, 0.15% formic acid) and eluted at a flow rate of 250 nL/min with a segmented 1-h gradient from 1 to 58% buffer B (80% acetonitrile, 0.15% formic acid). The data was acquired in data-independent acquisition (DIA) mode. The Orbitrap-based method used contained 40 dynamic windows over a scan range of 350–1650 m/z with 30,000 resolution and 27% HCD collision energy and a survey scan with 120,000 resolution and a maximum injection time of 50 ms and default charge state 2. The mass spectrometry proteomics data have been deposited to the ProteomeXchange [127] Consortium via the PRIDE [128] partner repository with the dataset identifier PXD049084. The processed protein information and protein-protein associations are available through the BioStudies repository for this paper https://www.ebi.ac.uk/biostudies/studies/S-BSST1423.

### Pre-processing of synaptosome co-fractionation data

Raw files were converted to HTRMS using HTRMSConverter (Biognosys) and analyzed in Spectronaut 13 (Biognosys) applying the “only protein group-specific” proteotypicity filter, and otherwise, the standard manufacturer’s settings for directDIA using the UniProt reference proteome for *Rattus norvegicus* (retrieved October 2019), QC standards and common laboratory contaminants. The Spectronaut output (BGS report) was then further pre-processed (Table S29). Rat protein identifiers were converted to human orthologs (Ensembl BioMart, date = 2023-09-26). The total sum of intensity of the fractions was scaled to account for different dilutions applied at the MS injection stage (fractions 44 to 54 were multiplied by 2 and fraction 56 to 75 were multiplied by 4). After rescaling, fractions were removed when the total sum of intensity was less than 50% of the average total sum of intensity for the adjacent 10 fractions (fractions 44, 45, 50, 51, 63 and 69). For the purpose of estimating co-fractionation through correlation of protein intensity across fractions, the total sum of intensity for each fraction was then normalized to account for variation of total intensity between neighboring fractions (∼25.8% difference in total intensity between neighboring fractions on average). The data was then pre-processed following ‘Computing interaction probabilities from co-fractionation data’.

### Predicting the synaptic interactome from co-fractionation data in the brain (Fig. 5a-b)

Co-fractionation data from the *in vivo* rat synaptosomes (75 fractions), *in vivo* mouse brains (110 fractions) [6] and subcellular fractionation of a human glioblastoma cell line (U251, 10 fractions) [7] was pre-processed as described in ‘Pre-processing of synaptosome co-fractionation data’ and following ‘Computing interaction probabilities from co-fractionation data’. For each study, the correlations of the pre-processed intensities were computed per protein pair across fractions as in ‘Computing association probabilities**’**. The correlation estimates of the studies were then filtered for protein pairs that were quantified for all three studies (1.287.210 pairs), and converted to interaction probabilities using a logistic model and all three studies as variates (model weights ∼0.90 (glioblastoma), ∼2.35 (mouse brain), ∼0.68 (rat synaptosome); intercept −0.95), with complex members as positives (following ‘Computing association probabilities’) (Table S30). AUC values for the recovery of complex members were computed using the association probabilities for each study individually and the combined studies following (‘Computing AUC values’).

### Interactions involving synaptic proteins enriched for more probable associations (Fig. 5c)

Sets of synaptic proteins were defined as reported in the SynGO database [83] (release date = 2021-02-25; 1233 genes), reported as synapse enriched in mouse brains [107] (converted to human proteins through orthologs - 1685 genes), reported as part of synaptic cellular components (genes in GO cellular components [88] whose name contain descriptions of ‘neuro’, ‘synap’, ‘axo’, or ‘dendrit’ components - 1869 genes), or reported as having elevated expression in the brain (Protein Atlas, maximum expression of the brain tissues exceeds twice the average expression of the other tissues - 6142 genes; [12]) (Table S31). Synaptic interactions were defined as the protein pairs whose interaction probability exceeded the 90-th percentile in the synaptic interactome (probability ∼0.67). Interactions were then split into involving synaptic proteins (as defined above) or between proteins that were not synaptic proteins (between other proteins). The distributions of association scores were then compared for the interactions involving synaptic proteins and the interactions between other proteins for each tissue in the association atlas and each set of synaptic proteins (one-sided MWU-test).

### Defining network of interactions between synaptic proteins (Fig. 5d)

The interaction probabilities of the synaptic interactome and the association scores of the brain from the association atlas were filtered for protein pairs that were quantified in both (1.287.206 protein pairs). The resulting protein pairs were further filtered for protein pairs that were both reported as synaptic proteins according to the SynGO database (41.616 pairs). Finally, the remaining protein pairs were filtered for both the synaptic interaction probabilities exceeding 0.5 and the brain association scores exceeding 0.5 (5.248 pairs). The resulting network of validated interactions between synaptic proteins was further annotated with the fractions of tissues for which the interactions were likely (from the association atlas) and whether the interactions were reported in the major protein interaction databases (STRING score > 400, HuMAP interactions exceeding the 99-th percentile, and interactions from HuRI, BioPlex, IntAct and CORUM complex members) (Table S14).

### Annotating validated synaptic interactions for brain disorders (Fig. 5d)

GWAS traits were selected for having associated mouse phenotype data (IMPC) or drug targets (ChEMBL - clinical stage II or higher). Traits were further filtered for having trait-level association scores relative to the median association score in each tissue being elevated in the brain compared to the other tissues (z-score > 0 - 43 traits; 11 disorders were specific to the brain; Table S15). The network of synaptic interactions was filtered for interactions between proteins associated with any of the brain-specific disorders, and each interaction was annotated with the respective disorder(s) (72 proteins; 514 interactions) (Table S16). Finally, the network was split into subnetworks of synaptic interactions between proteins associated to each disorder - if at least three interactions were available - and further annotated with evidence for that disorder (whether the gene is associated to the disorder according to IMPC, ChEMBL, or OTAR GWAS, including the SNPs and L2G scores for each gene). The visualized networks represent the annotated data (except for schizophrenia).

### List of materials

| Cell lysis + SEC + sample preparation |  |  |
| --- | --- | --- |
| # compound | # product number | # supplier |
| HEPES, BioPerformance ≥ 99.5 | H4034 | Sigma-Aldrich |
| Sodium-Chloride (NaCl) | 1.06404.5000_P | Merck |
| Sodium-Fluoride (NaF) | S7920 | Sigma-Aldrich |
| Sodium Orthovanadate (Na<sub>3</sub>VO<sub>4</sub>) | S6506 | Sigma-Aldrich |
| PMSF | P7625 | Sigma |
| Protease Inhibitor cocktail | P8849 | Sigma |
| n-Dodecyl β-D-maltoside (DDM) | D4641 | Sigma-Aldrich |
| Column Performance Check Std. Aqueous SEC 1 | AL0-3042 | Phenomenex |
| BCA assay kit | 23225 | ThermoFisher Scientific |
| Ultracentrifuge tubes (8×34 mm) | 343776 | Beckman-Coulter |
| TLA 100.1 rotor |  | Beckman-Coulter |
| Amicon(R) Ultra-4, 30 K | UFC803096 | Merck |
| Guard column, SRT-C1000, 5 μm, 1000 Å, 7.8×50 mm | 235950-7805 | Sepax |
| SRT-C1000, 5 µm, 1000 ,<br>7.8×300 mm | 215950-7830 | Sepax |
| Liquid Chromatography high<br>pressure pump | 1100 Series | Agilent |
| Degasser | 1100 Series | Agilent |
| LC Fraction Collectors | 1260 Infinity Series | Agilent |
| LC Operating Software<br>Agilent |  | OpenLab |
| AcroPrep Advance 96 Well<br>Filter Plates, 350 µL,<br>Omega 10K MWCO | 8164 | Pall |
| HPLC-grade H2O | 7732-18-5 | Fisher Chemical |
| Urea | GEPURE00-67 | Eurobio |
| TCEP | C4706 | Sigma-Aldrich |
| Iodoacetamide | I6125 | Sigma |
| Ammonium-bi-carbonate<br>(Ambic) | 09830 | Fluka |
| Sequencing Grade Modified<br>Trypsin | V5113 | Promega AG |
| Lysyl Endopeptidase, Mass<br>Spect. Grade | WA3 125-05061 | Wako |
| Acetonitrile, LC/MS Grade | 75-05-8 | Fisher Chemical |
| LoBind tubes | 022431081 | Eppendorf |
| iRT Kit | Ki-3002-2 | Biognosys |

## SUPPLEMENTARY FIGURES

**Figure S1.**
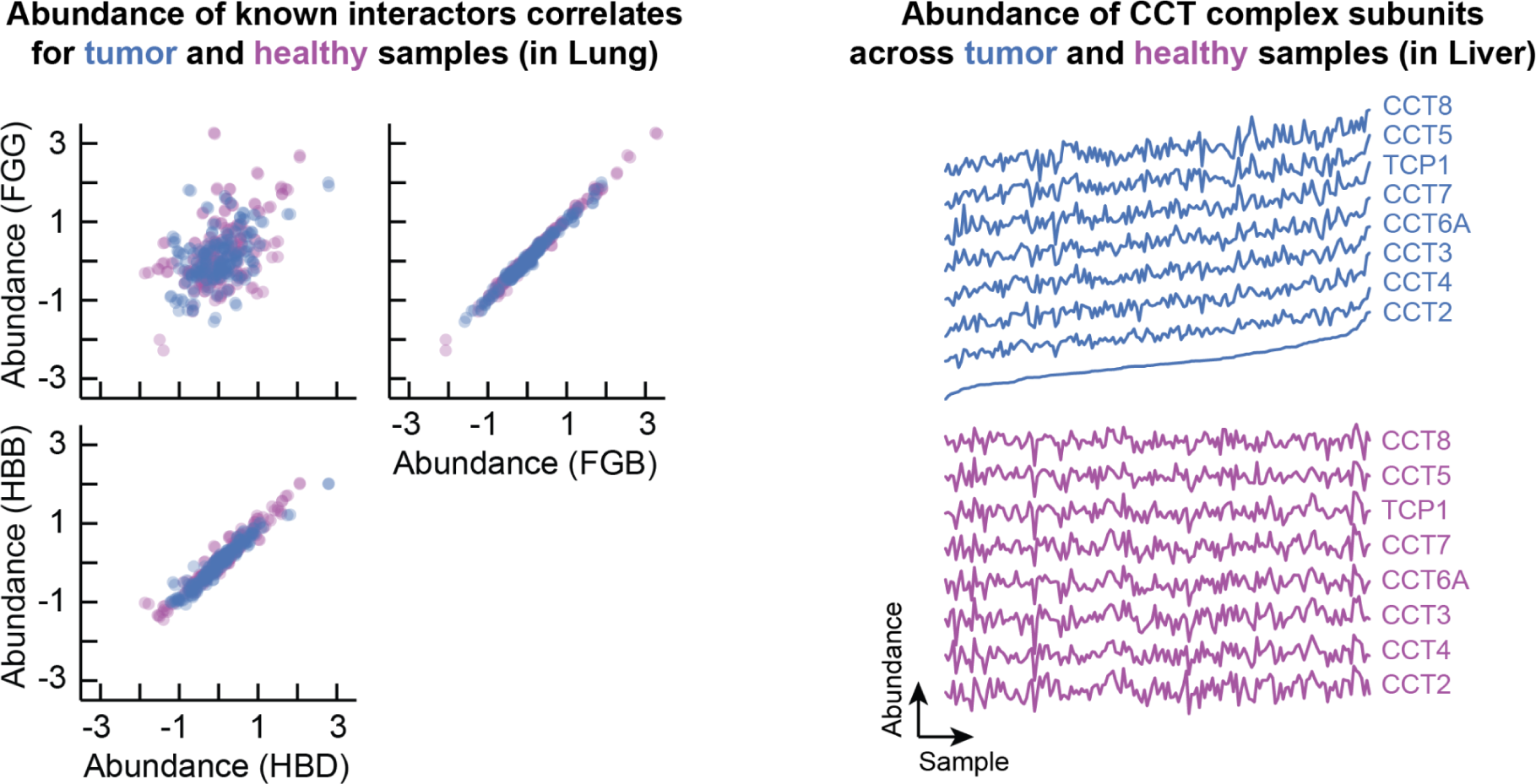
Abundance of known interactors correlates across tumor and healthy samples (Related to Fig. 1). **(a)** To compare the abundance of proteins across samples, we filtered for studies for which we had both tumor and healthy samples whose abundance values were not truncated (17 studies). We further filtered each study to only consider paired samples and genes that were quantified in all samples for both the tumor and healthy samples. As an example of the co-abundance of known protein interactions, we looked at the protein pairs that were most co-abundant in tumors on average across studies. Specifically, we took the Fibrinogen proteins FGB and FGG (average Pearson correlation ∼0.97) and the Hemoglobin proteins HBB and HBD (∼0.93). Shown is the (co-)abundance of these proteins for the healthy and tumor samples for a representative cohort with many samples (n = 216 - in the lung). Each dot represents the abundance in one sample for the healthy (purple) and tumor (blue) samples. **(b)** As an example of the co-abundance of subunits in a protein complex, we selected the abundance of the CCT complex subunits from a representative cohort having a large number paired tumor and healthy samples (n = 163 - in the liver; Gao2019). Shown is the abundance of the CCT subunits across patients for the tumor (blue - top) and paired healthy samples (purple - bottom). Patients were sorted by the abundance in tumors of the subunit that had the largest variance across patients. Subunits were sorted by the correlation across tumors with the same subunit (CCT2). Healthy data is shown in the same order. For this example, we found that the abundance of CCT subunits correlates across tumor samples (average Pearson correlation coefficient ∼0.93 ± 0.01) and healthy samples (∼0.75 ± 0.01). Surprisingly, we found that the abundance of CCT subunits poorly correlated between the healthy and tumor dissections across patients (0.11 ± 0.04), which generalized to the correlation of protein abundance of paired healthy and tumor samples across studies (average correlation coefficient of healthy and tumor samples across common genes and studies ∼0.21 ± 0.03).

**Figure S2.**
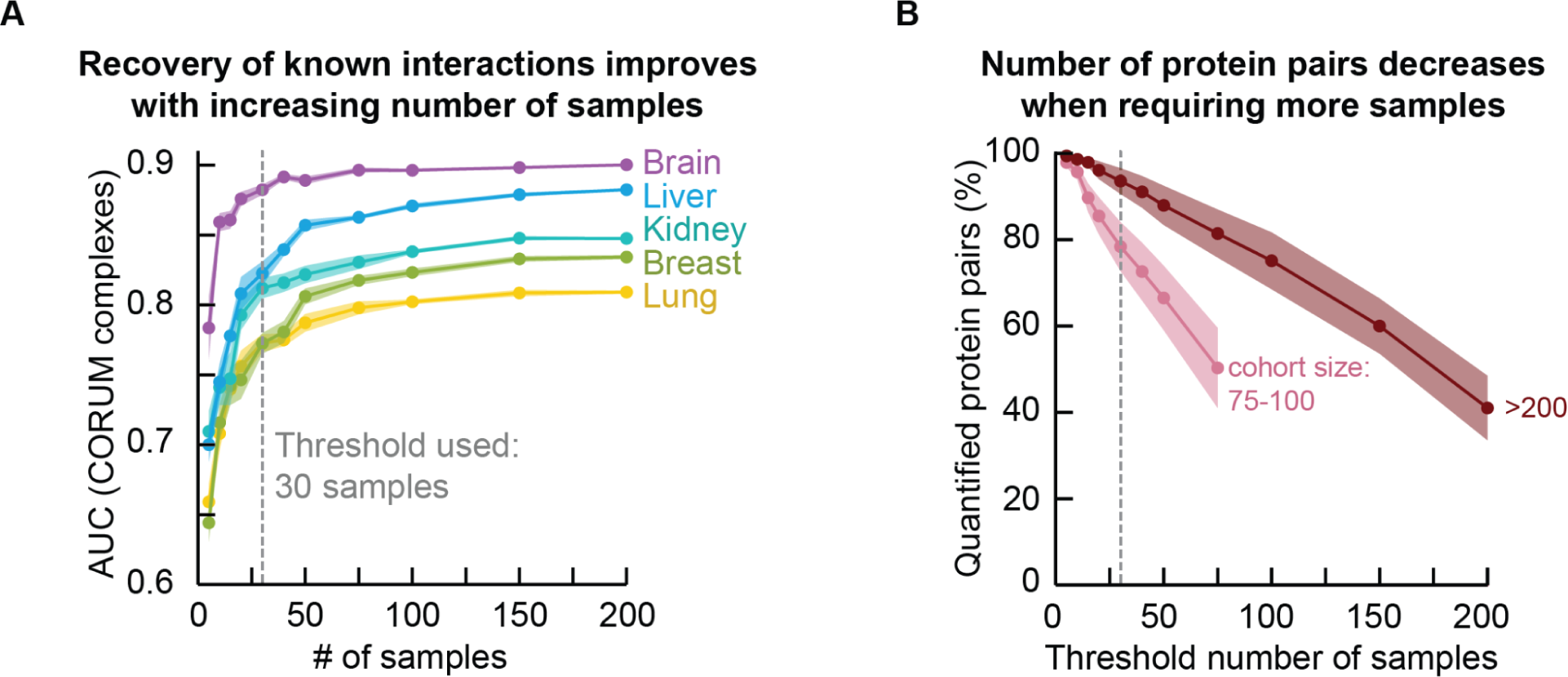
Co-abundance estimates as function of the number of patients per cohort (Related to Fig. 1). **(a)** To test the effect of the available number of samples on the co-abundance estimates, we computed the recovery of known protein interactions when varying the number of samples. Specifically, we computed the AUC for the recovery of interactions between subunits of the CORUM complexes when subsampling the samples of cohorts with large numbers of samples. As cohorts, we selected studies that had cohorts consisting of at least 200 cancer patients and whose AUC on the entire dataset exceeded 0.75. Moreover, we used at most one cohort per tissue, selecting the cohort achieving the highest AUC value on the entire dataset. In total, we selected 5 cohorts (breast, kidney, liver, lung and brain). We then filtered each cohort for proteins that were quantified for all samples, such that we could quantify co-abundance estimates for the same protein pairs across sample sizes (i.e., the analysis does not depend on the available protein pairs). We then subsampled varying numbers of patients of each cohort and computed the AUC for the recovery of interactions for CORUM complexes using the association probabilities computed from the co-abundance of proteins within the respective subsampled sets of patients. Shown is the AUC as a function of the number of samples for each of the cohorts. As expected, the AUC value decreases as the number of samples decreases. **(b)** Additionally, we computed the number of protein pairs that were both quantified in at least a threshold number of samples as a function of that threshold number. Here, we did not filter for proteins quantified for all samples. Shown is the percentage of possible protein pairs as a function of the threshold number of samples required for computing a co-abundance estimate (same studies as in (a) (dark red), studies with 75-100 samples (light red)). The fraction of protein pairs for which we can compute a co-abundance estimate decreases when requiring more samples (increasing threshold number of samples). In (a-b), the gray dotted line shows the threshold number of samples (n = 30) used throughout this work. Overall, these analyses demonstrate that having more samples for computing co-abundance estimates result in association scores for less protein pairs but higher AUC values. As a compromise, we used a threshold of 30 samples for computing co-abundance estimates throughout all analysis.

**Fig. S3.**
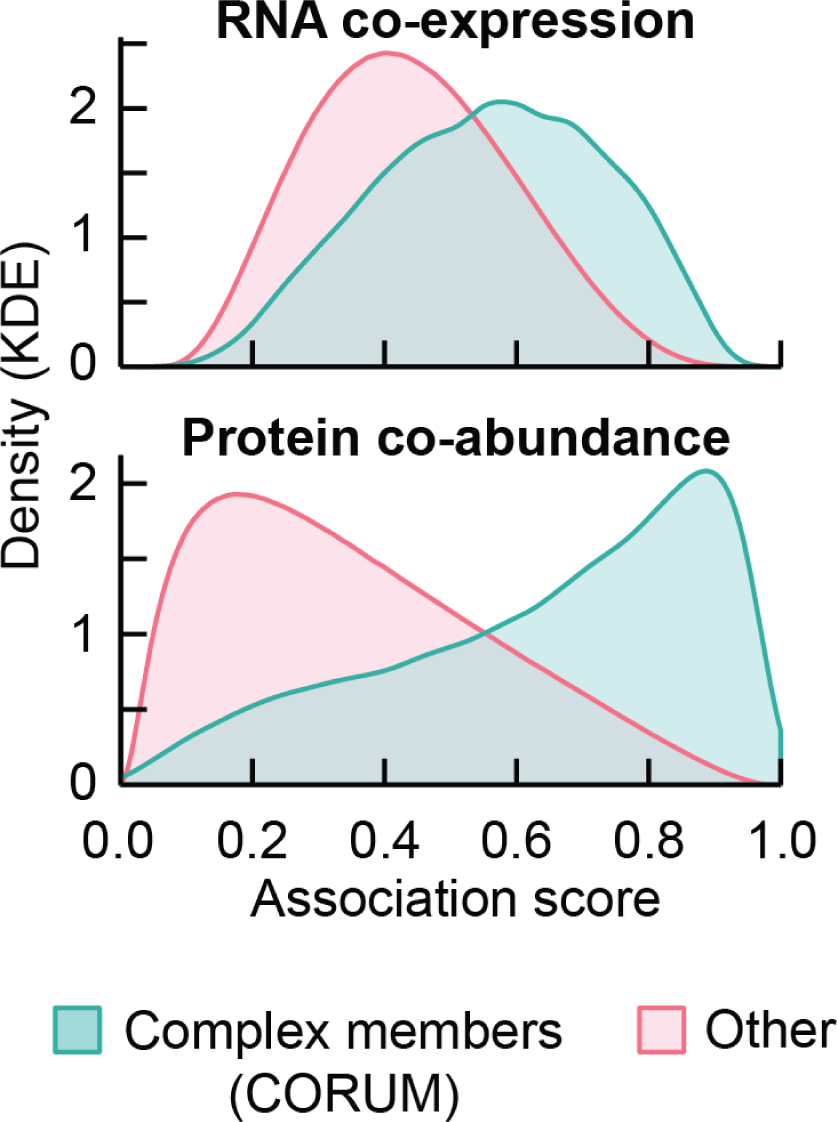
Associations between complex members are more likely than associations of other protein pairs (Related to Fig. 1). We compared the association scores of complex members with the association scores of proteins that are not reported as interacting according to the CORUM database using the transcriptomics and proteomics data from a single Lung cohort. Shown are the Kernel Density Estimates (KDE) for the association scores derived from protein co-abundance and RNA co-expression, for both complex members (green) and other protein pairs (red). These distributions demonstrate that the associations between complex members are more likely compared to the associations of other proteins, especially as quantified through protein co-abundance.

**Figure S4.**
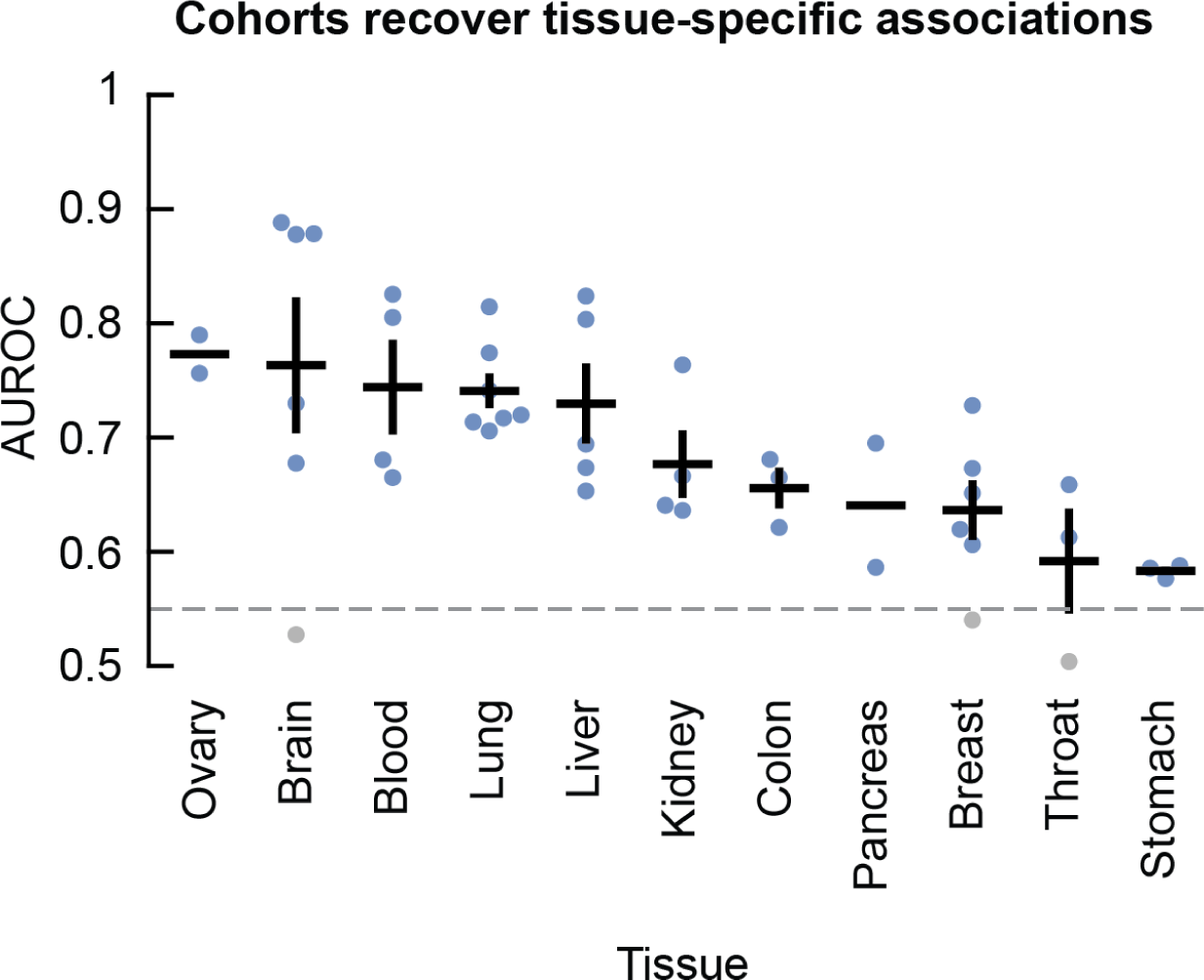
Cohorts recover tissue-specific associations (Related to Fig. 1). We tested whether a cohort of a tissue could recover associations predicted by the other replicate cohorts of the same tissue. For this, we used the tissue-specific associations for each tissue: associations whose average probability exceeded the 95-th percentile for a given tissue and whose average probability remained below 0.5 across all other tissues. We then used a hold-one-out methodology where we predicted the tissue-specific associations for a given tissue with all-but-one cohorts of that tissue, and tested how well the withheld cohort could recover the predicted associations. By predicting and recovering the tissue-specific associations through sequentially withholding each cohort of a given tissue, we could score how well each cohort aligned with the other cohorts of that tissue. Shown are the AUCs for recovering the tissue-specific associations across tissues. Each dot represents one cohort. Error bars show mean and s.e.m. (when available). We found three cohorts whose AUC values were below 0.55 for recovering the associations predicted by the other cohorts of the same tissue. We chose to not incorporate these cohorts for computing the aggregated tissue scores as the tissue-specific associations of these cohorts were poorly aligned with the other cohorts of the same tissue (brain, breast and throat).

**Figure S5.**
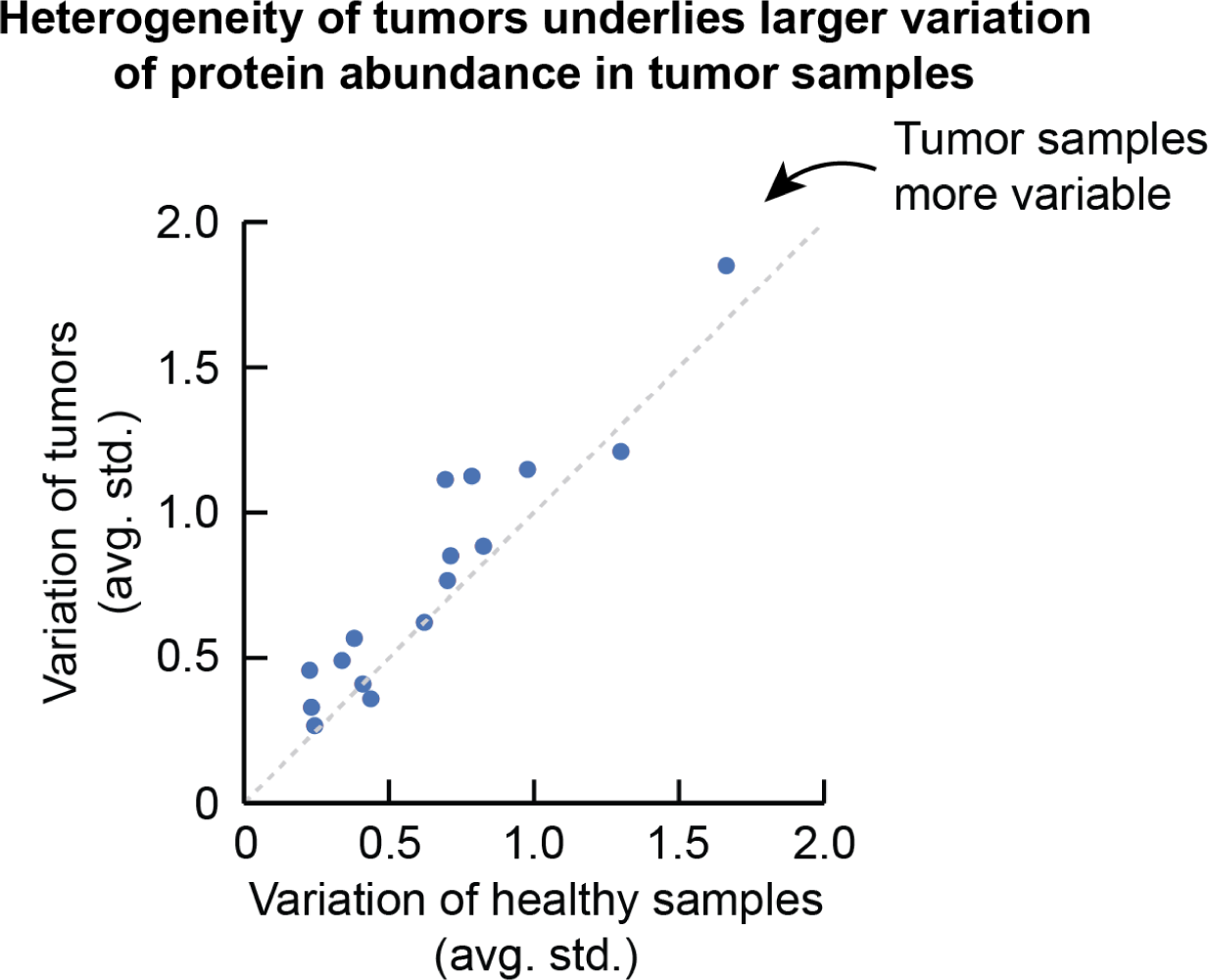
Comparing variation of abundance for tumor and healthy samples (Related to Fig. 2). To compare the variation between samples of tumor and healthy samples, as before, we selected the abundance of common genes and common samples in the 17 studies that had paired tumor and healthy samples available. We then computed the standard deviation of abundances across samples and averaged the standard deviations over all genes. We found that the average standard deviation across patients was 0.78 ± 0.1 for the tumor samples and 0.66 ± 0.1 for the paired healthy samples. Shown is the variation across tumor samples as function as the variation across healthy samples. We compared the distributions of average standard deviations using a paired t-test (one-sided; p-value 1.9e-3), suggesting that - averaged over all genes - there is no reason to assume that the tumor samples are less variable than the paired healthy samples. Finally, when comparing the standard deviation of abundances across samples between tumor and healthy samples per gene, we found that the tumor samples were on 1.3 ± 0.08 -fold more variable compared to the healthy samples. Overall, these analyses suggest that the genetic heterogeneity of tumors is represented in the data by a larger variation in abundance of the tumor samples compared to the paired healthy samples in the cohorts. Dots represent the different studies. Diagonal indicates equal variation between tumor and healthy samples.

**Fig. S6.**
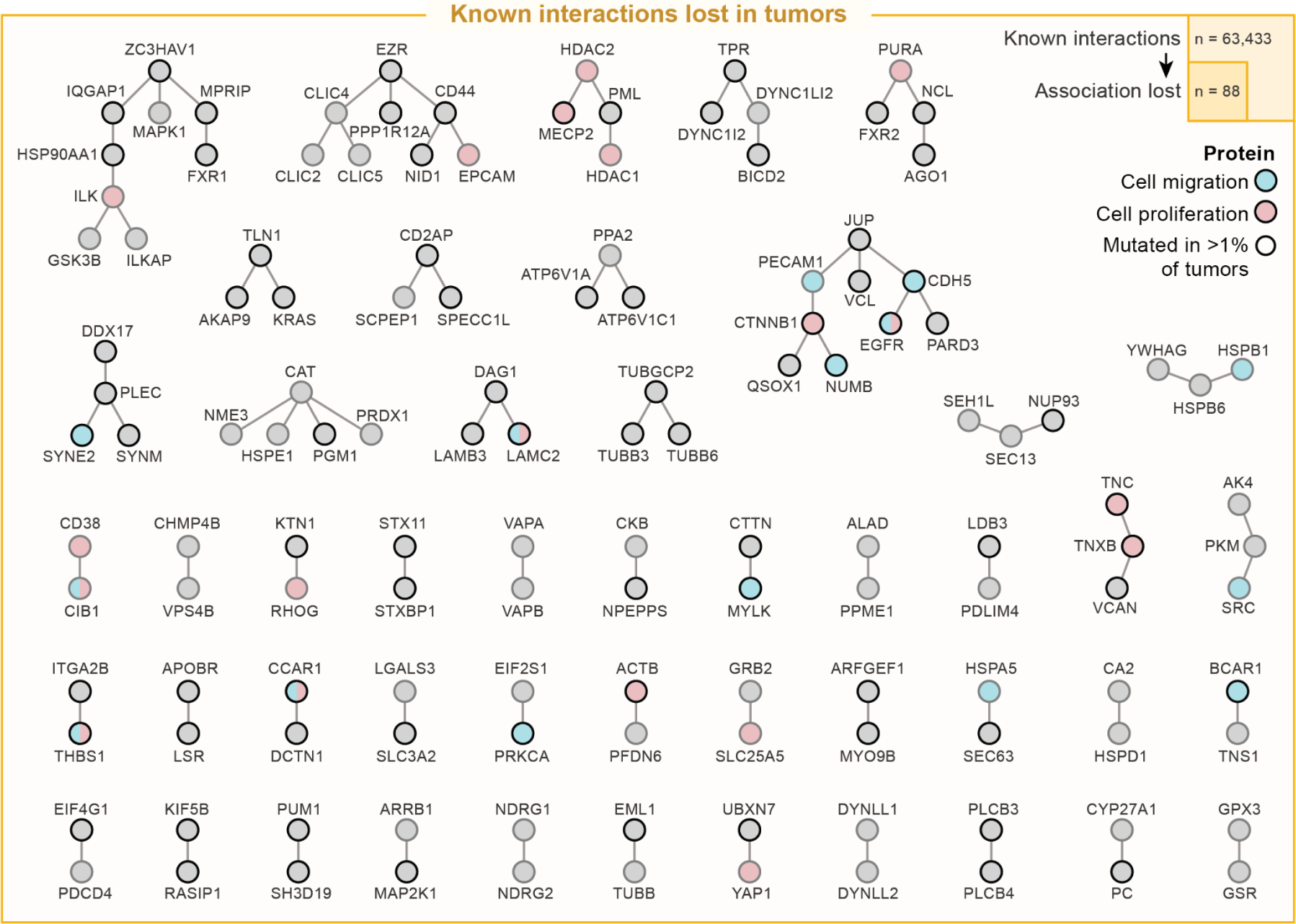
Network of protein interactions that are lost in tumors (Related to Fig. 2). For six tissues we had protein association scores derived from both tumor and from healthy biopsies (colon, kidney, liver, lung, stomach, throat). We hypothesized that there may be some protein interactions that are lost for tumors and that the loss of these interactions could be reflected in our association scores. To test this, we computed the difference between the healthy and tumor-derived association scores for each of the six tissues. We then selected the protein pairs that were quantified in at least three of the tissues (13,905,554 protein pairs). We further filtered for protein pairs that are known to interact (reported in STRING with scores exceeding 400 – 63,433 pairs). Finally, we selected the protein pairs for which, in at least one tissue, the tumor-derived association score was at least 0.5 below the association score derived from the healthy biopsies of the same tissue (“lost associations” – 88 pairs). Thus, we ensured that we considered only known interactions that were commonly quantified in our association atlas and whose likelihood of associating substantially decreases for at least one tissue. We found no reason to assume that known protein interactions were enriched for such lost associations compared to associations without prior evidence (i.e., no STRING score exceeding 400; p-value 0.7 – one-sided Fisher exact test). Here, we used all non-“lost associations” with and without having prior evidence as backgrounds respectively. Next, we explored the proteins that were involved with the lost associations. First, using mutation frequencies for any tissue from the Pan-cancer Atlas (cbioportal), we found that 58% of the proteins with lost associations were frequently mutated in cancers (mutation frequencies exceeding 1%). Indeed, the average decrease in association scores from healthy- to tumor-derived scores was generally larger for cancer proteins compared to the association scores between other proteins (p-value 6.7e-207 – one-sided MWU-test). Looking at the GO biological processes related to the lost protein associations, we found that the lost associations were most enriched for proteins related to positive regulation of cell population proliferation (p-value = 6.4e-5; GO:0008284) and positive regulation of cell migration (2.5e-3; GO:0030335), with fibroblast migration (1.5e-2; GO:0010761), canonical Wnt signaling (1.5e-2; GO:0060070) and adherens junction organization (1.5e-2; GO:0034332) also being amongst the most related processes (Fisher’s exact test with Benjamini-Hochberg (BH) adjusted p-values; all other proteins from the known interactions and all other proteins in GO biological processes used as backgrounds respectively). Finally, we found that 0.027% p/m 0.02% of likely associations had a difference that exceeded 0.5 when comparing the healthy and tumor-derived scores. In other words, at most 0.03% of protein pairs have a substantial difference in association scores and are likely to be associating according to either – but not both – the healthy-derived or the tumor-derived scores. The substantially different association scores are thus negligible on the scale of the association atlas. Thus, while the known interaction lost in tumors may be sensible in the context of cancer, our observations demonstrate the lack of a sizable cancer-specificity in the association atlas. The visualized network shows the known protein interactions whose association is lost in tumors for at least one tissue. Nodes are colored for the proteins being associated with the positive regulation of cell population proliferation (red), cell migration (blue) or other (gray). Node edges indicate proteins that are frequently mutated in cancer (black) or not (gray).

**Figure S7.**
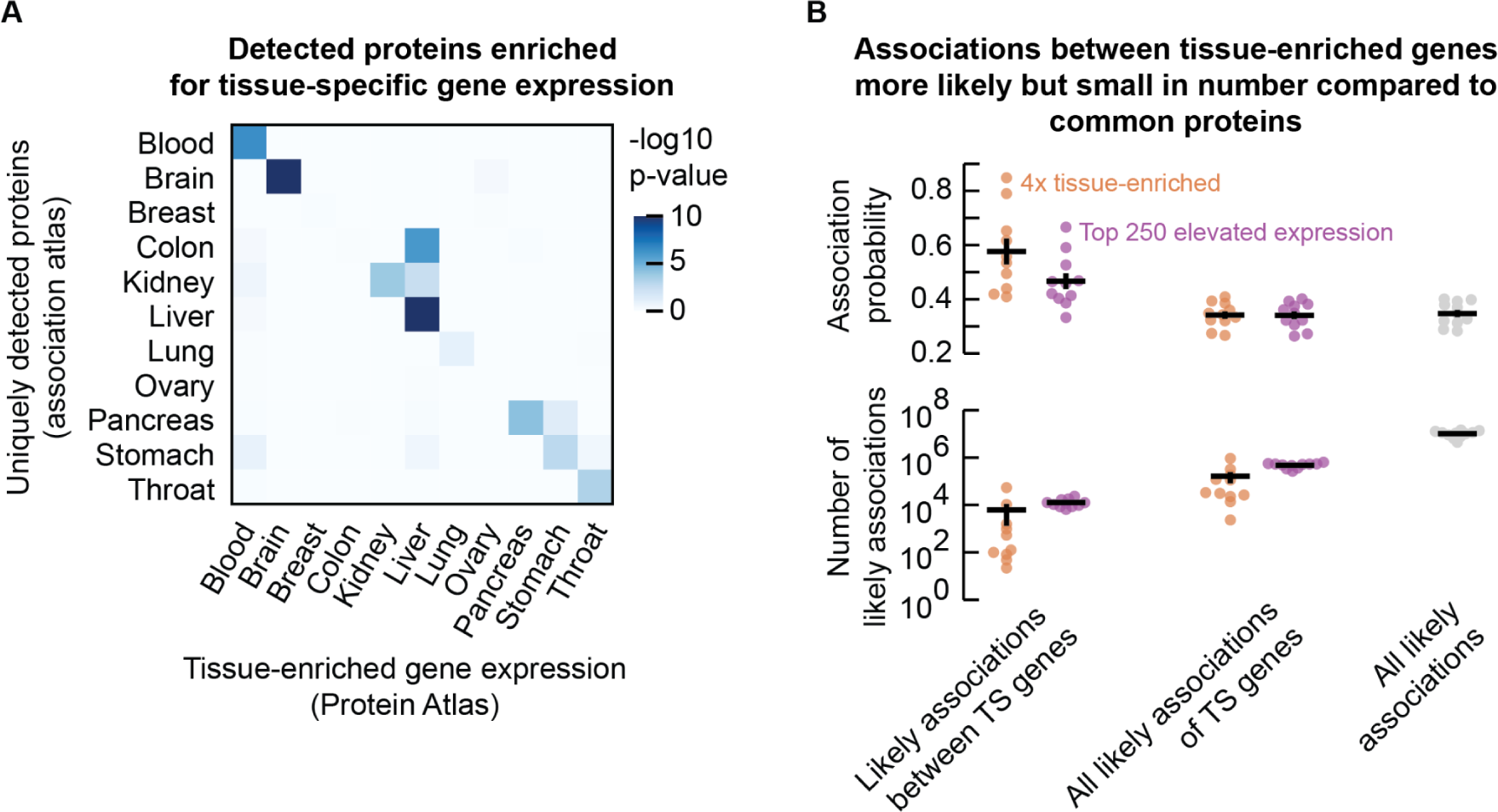
Differences between tissues not primarily driven by gene expression (Related to Fig. 2). **(a)** Heatmap showing the enrichment (log10 p-value; Fisher’s exact test) of genes with tissue-enriched expression (at least 4-fold higher expression compared to any other tissue; from the Protein Atlas ‘consensus normalized expression’) amongst the proteins for which we quantified protein associations in each tissue. **(b)** association probability and number of likely associations of proteins whose expression is tissue-specific (“TS”). Shown is the median association probability in each tissue for all quantified protein pairs (gray dots), all associations of the tissue-specific proteins, or the associations between the tissue-specific proteins. As tissue-specific proteins, we used the definition of the Protein Atlas (“tissue-enriched”; genes whose expression is at least 4-fold higher in one tissue compared to any other tissue – orange dots) or the top 250 proteins that are quantified in our association atlas and had the most tissue-specific expression (“elevated expression”; highest z-score across tissues – purple dots). Each dot represents one tissue. Given the low and varying number of proteins that were tissue-enriched for some tissues, we used here the top 250 proteins with elevated expression as representative to quantify the effect of gene expression. We found that the median association probabilities changed from 0.35 ± 0.01 for all quantified protein pairs to 0.34 ± 0.01 for all associations of proteins with elevated expression, and to 0.47 ± 0.03 for the associations between the proteins with elevated expression. Here, the likely associations between the proteins with elevated expression represented 0.14% ± 0.02% of all likely associations, while all likely associations of the proteins with elevated expression represented 4.8% ± 0.2% of all likely associations of the tissues. We found similar numbers for the tissue-enriched genes (main text). Thus, the proteins that were quantified for a given tissue are generally enriched for genes with elevated expression for that same tissue, but not the other tissues. Moreover, the protein associations between these genes are more likely than the associations of other proteins. However, the associations of proteins with elevated expression are not generally more likely compared to the associations of all proteins, and the collection of likely associations of proteins with elevated expression is not substantial compared to all likely associations.

**Figure S8.**
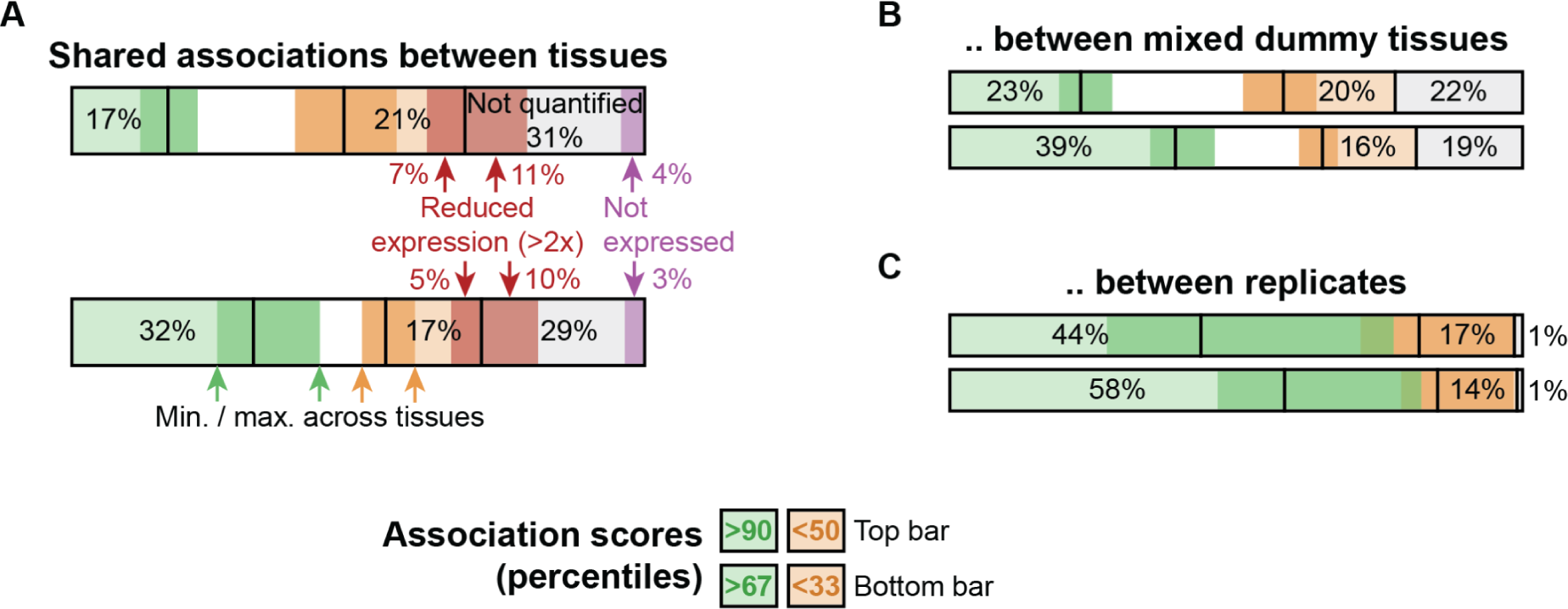
Quantifying differences of associations between pairs of tissues (Related to Fig. 2). **(a)** For each tissue, we selected the protein pairs whose association scores exceeded the n-th percentile of associations in that tissue (“likely” associations - for convenience of notation, we changed the definition for likely associations to be the associations exceeding some n-th percentile of associations in this supplementary figure). We then quantified the percentage of protein pairs whose association score in another tissue also exceeded the n-th percentile (green bars), remained below some m-th percentile (“unlikely” - orange bars), or that were not quantified in the other tissue (“not quantified” - gray bars). Shown is the percentage of associations averaged over all pairs of tissues. Dark colored sections (green/orange) indicate the maximum and minimum percentage of the tissue-level averages. For the unlikely and unquantified associations we additionally computed the percentage of associations that involved proteins whose expression was reduced by at least 2-fold in the other tissue (red bar section - Protein Atlas) or that were not expressed (purple bar section). Shown are the percentages for unlikely associations as defined through association scores exceeding the 90-th percentile (top bar) or 67-th percentile (bottom bar), with the unlikely associations defined by association scores remaining below the 50-th percentile (top bar) or 33-rd percentile (bottom bar). We find that 17% - 32% of associations are shared with other tissues depending on the threshold used to define the likely associations. Moreover, similar to Fig. 2e, we find that 18% - 22% of differences in likely associations between tissues are explainable by changes in gene expression. With a wide range of thresholds defining the likely and unlikely associations, these observations support our observation that the differences between tissues are not driven by gene expression. **(b-c)** As in (a) but for associations shared between “dummy tissues” (b) and the healthy- and tumor-derived scores as independent replicates for the same tissue (c). **(b)** Analogously to the 11 human tissues, we constructed “dummy tissues” that aggregated the association probabilities from cohorts of mixed tissues. By mixing cohorts of different tissues, we eliminate the tissue-specificity of tissues (i.e., the effects of tissue-specific associations and gene expression), thereby leaving other sources of variation. That is, the difference between dummy tissues and actual tissues is primarily the tissue-specificity of our association atlas. Here, we mixed cohorts originating from the lung, breast and liver (the tissues having at least 5 cohorts, excluding the brain). Having 5 cohorts available from each, we could construct 25 dummy tissues where all cohorts are paired exactly once (25 unique combinations of cohorts from the breast and liver, that each can get assigned a lung cohort to never reuse a pair of cohorts). We then split these dummy tissues into 5 sets of 5 dummy tissues each, ensuring that in each set the dummy tissues do not share any cohorts. For each set, we then computed the differences of associations between pairs of dummy tissues (as in (a)). We found that - depending on the thresholds for likely and unlikely associations - between 23% and 39% of likely associations was shared between dummy tissues. Compared to our association atlas, the dummy tissues thus share 6% - 7% more likely associations, which can be attributed to tissue-specificity of our association atlas. Given the conservation of other sources of variation for the dummy tissues, these results provide a lower bound of 6% for the percentage of tissue-specific associations between pairs of tissues. Moreover, the unquantified associations for the dummy tissues illustrate that 19% - 22% of likely associations are not replicated when using different cohorts and studies for the same tissue. This observation suggests that the majority of the unquantified associations between tissues in the association atlas originated from experimental sources such as the different groups of patients and experimental procedures, further confirming our observations that the differences between tissues are not driven by gene expression. **(c)** As in Fig. 2e for the healthy- and tumor-derived scores. We find that between 44% and 58% of associations are shared between replicate tissues, resulting in an estimated 26-27% of associations being tissue-specific when compared to the association atlas. Together, these results suggest that - for the chosen thresholds for likely and unlikely associations - between 6% and 27% of likely associations are tissue specific between pairs of tissues of our association atlas.

**Figure S9.**
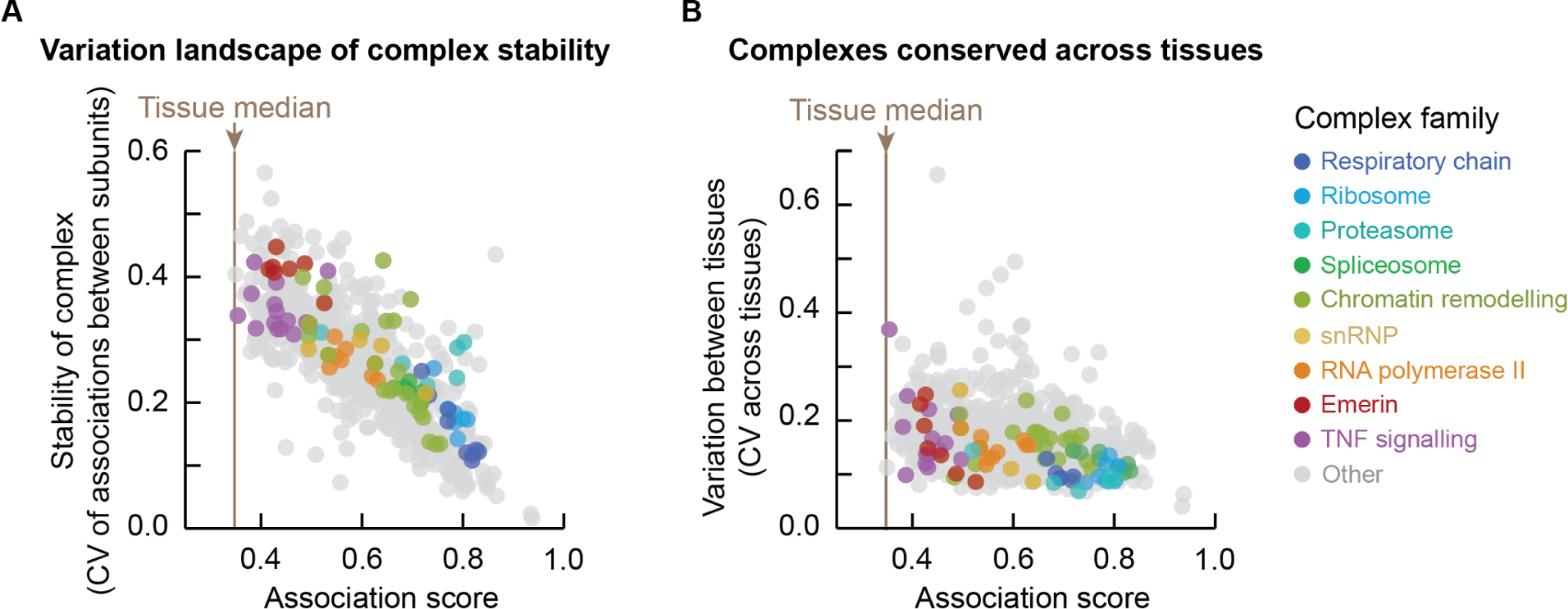
Well-defined human protein complexes preserved across tissues (Related to Fig. 2). Given the conservation of associations between complex members across tissues, we explored the variation landscape of well-defined human protein complexes. Specifically, we filtered the human protein complexes in the CORUM database for complexes that consisted of at least 5 subunits and merged complexes with identical names. For the resulting 645 protein complexes, we explored the variation of associations within complexes and between tissues. We computed a complex-level association score (the median association score between all pairs of subunits; Table S17), a complex-level variation (the coefficient of variation (CV) of the association scores of all pairs of subunits; Table S18) and the variation of complexes between tissues (the CV of the complex-level association scores across tissues). **(a)** Complex-level variation as a function of the complex-level association scores, both averaged across tissues. We found that 76% of the protein complexes had an average complex-level association score that exceeded 0.5. Complexes became more variable as the complex-level association score decreased (rho = −0.77, −log10-p-val = 125), suggesting that the associations of subunits become unlikely for more variable – less stable – complexes. Indeed, we found that variable complexes typically involved signaling and regulation (e.g., TNF and Emerin complexes) while more stable complexes involved central cellular structures (e.g., ribosomes and the respiratory chain). **(b)** Average and variation of complex-level association scores across tissues. Variation of complexes between tissues poorly correlates with the complex-level association scores (rho = −0.27, p-value = 2.8e-12). As expected, central cellular structures such as the ribosome, respiratory chain, spliceosome, proteasome, and RNA polymerase were less variable between tissues compared to other protein complexes for which we quantified at least 5 subunits in all tissues (average CV across tissues of ∼0.12 and ∼0.15 respectively, p-value 4.1e-7; MWU-test). However, we also found that complexes were not more variable across tissues than arbitrary collections of associations. Specifically, we found that there was no reason to assume that the complexes were more variable across tissues than dummy complexes consisting of arbitrary subunits and following the same size distribution (average coefficients of variation of 0.167 and 0.168 respectively; p-value 0.52, MWU-test. To construct these dummy complexes, we randomly sampled the subunits for each CORUM complex from the pool of proteins that was quantified in all tissues (20 replicates) and subsampled these sampled subunits to match the number of actual subunits that were quantified for each tissue. We then computed the association score of the dummy complex as the median association score between all pairs of the sampled subunits). Together, these observations suggest that the well-defined protein complexes are generally preserved across tissues, and that there is no reason to assume tissue-specificity at the complex-level beyond what would be expected to result from tissue-specific interactions of the individual subunits to a similar degree as arbitrary pairs of proteins. In (a-b), each dot represents one protein complex. Brown solid line represents the median tissue score averaged over the tissues.

**Figure S10.**
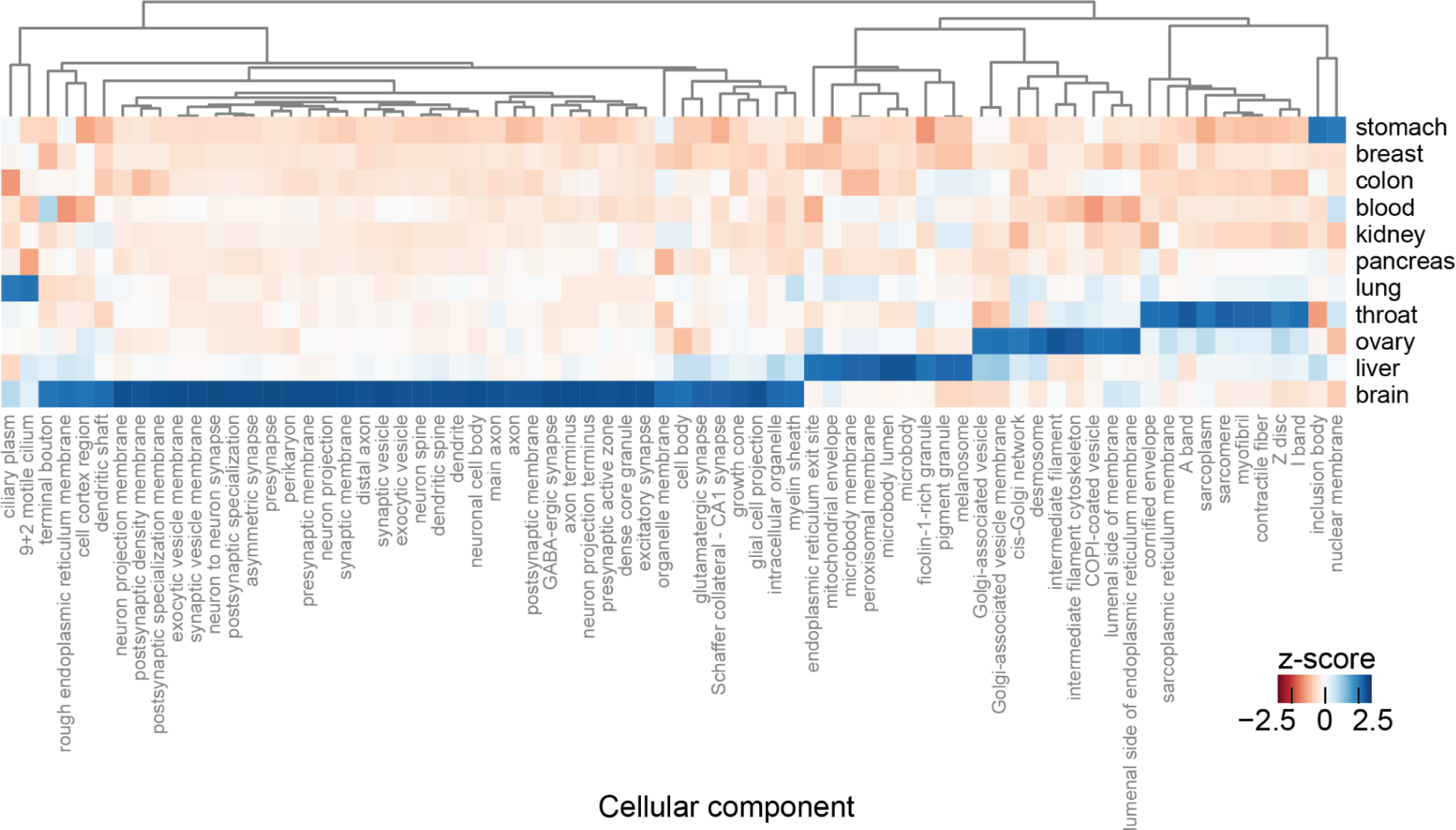
Protein associations vary for tissue-specific cellular components (Related to Fig. 2). Analogous to the protein complexes, for each tissue, we computed component-level association scores as the median association score between all pairs of proteins associated with cellular components as defined by the Gene Ontology (Methods). We then selected the cellular components for which at least 10 associated proteins were quantified in all tissues, and computed the component-level association scores relative to the median association score for each tissue. Shown are these relative component-level associations z-scored across tissues, filtered for the cellular components whose z-score exceeded 2.5 in at least one tissue. We found that ∼21% of cellular components had z-scores exceeding 2.5 in at least one tissue. Indeed, the protein associations varied strongly for typical tissue-specific components, such as synaptic components for the brain and structural components of muscle fiber for the throat.

**Figure S11.**
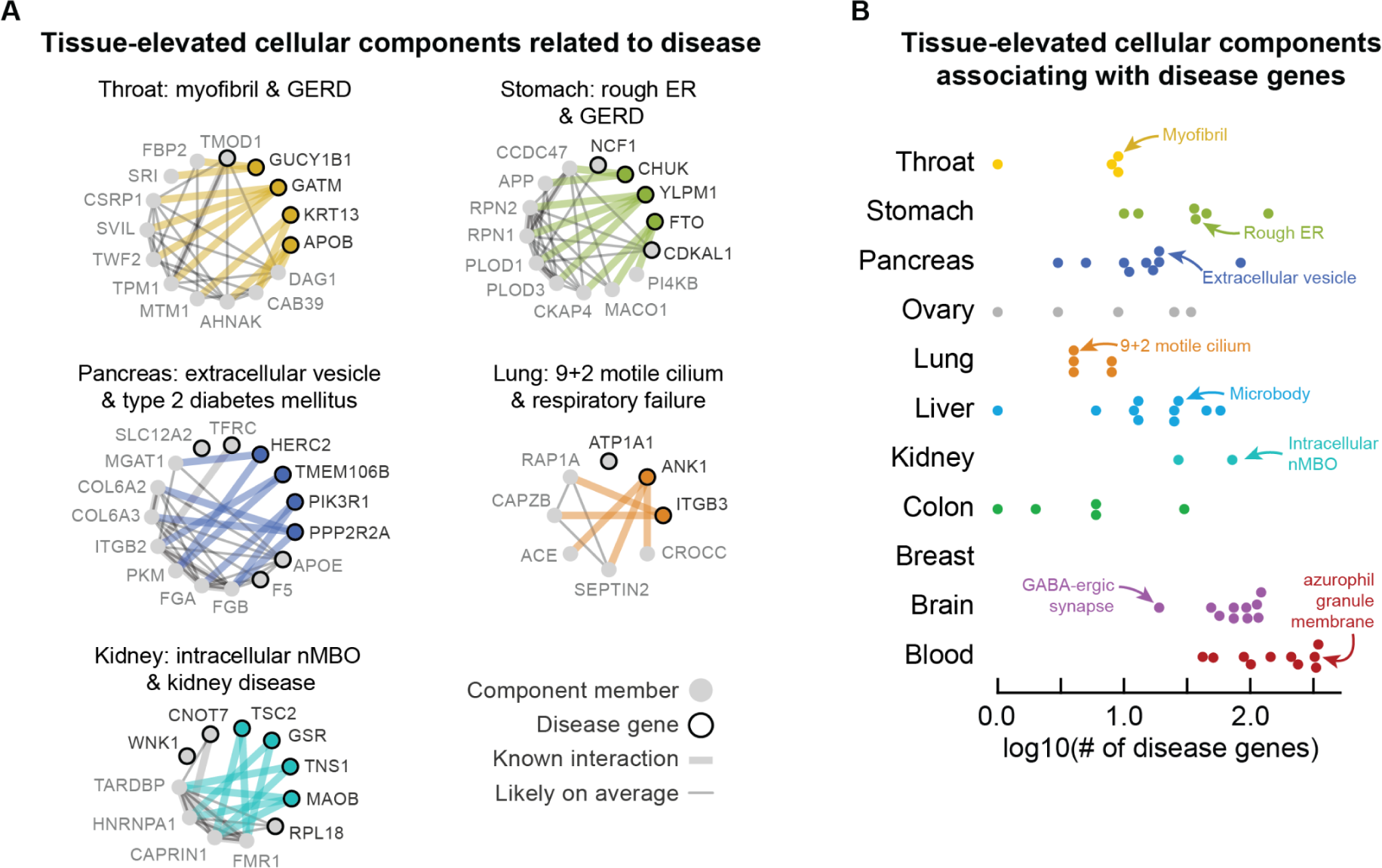
Tissue-specific associations for cellular components related to tissue-specific disorders (Related to Fig. 3). Analogous to Fig. 3b. We selected the cellular components for which at least 5 associated proteins were quantified in all tissues. For each tissue, we then filtered for the components whose relative component-level association score exceeded 1 when z-scored across tissues. For each tissue-elevated component, we selected tissue-related disorders whose associated proteins (partially) overlapped with the component, and looked for tissue-specific associations between the non-overlapping proteins (i.e., tissue-specific associations of proteins associated to the component with proteins associated to the disorder). Here, we defined as tissue-specific the associations that are likely in a given tissue (association score > 0.5) but not likely in any other tissue (< 0.5), and whose z-scored association score additionally exceeded 2 for that tissue. Finally, we filtered for proteins that had at least two of such tissue-specific associations for a given component-disorder pair. **(a)** Throat-specific associations of myofibril with proteins associated to Gastroesophageal Reflux Disease (GERD), stomach-specific associations of rough Endoplasmic Reticulum (ER) with GERD, pancreas-specific associations of extracellular vesicles with type 2 diabetes, lung-specific associations of 9+2 motile cilium with respiratory failure, and kidney-specific associations of intracellular non-Membrane-Bounded Organelles (nMBO) and kidney disease. Associations are likely on average across tissues (thin gray lines) or tissue-specific (thick colored lines). Thick gray lines are interactions with prior evidence (STRING scores exceeding 100). Shown are component members (gray nodes) that are disease genes (black edge) or have tissue-specific associations with disease genes (colored nodes). **(b)** Disease-genes having tissue-specific associations with tissue-elevated cellular components. Shown is the number of disease-associated proteins having at least 2 tissue-specific with each tissue-elevated component (overlapping genes are ignored). Each dot represents one tissue-elevated component (no components were elevated for the breast). In (a-b), we ignored traits (measurements) and cancers, and required disorders to be related to tissues by name (e.g., disorders for hemoglobin and anemia were assigned as related to blood, while “neuro” disorders were related to the brain, etc). Proteins were associated with disorders through GWAS (OTAR L2G score > 0.5), mouse phenotyping (IMPC) or as drug targets (ChEMBL clinical stage II or higher). Components are defined through GO annotation.

**Figure S12.**
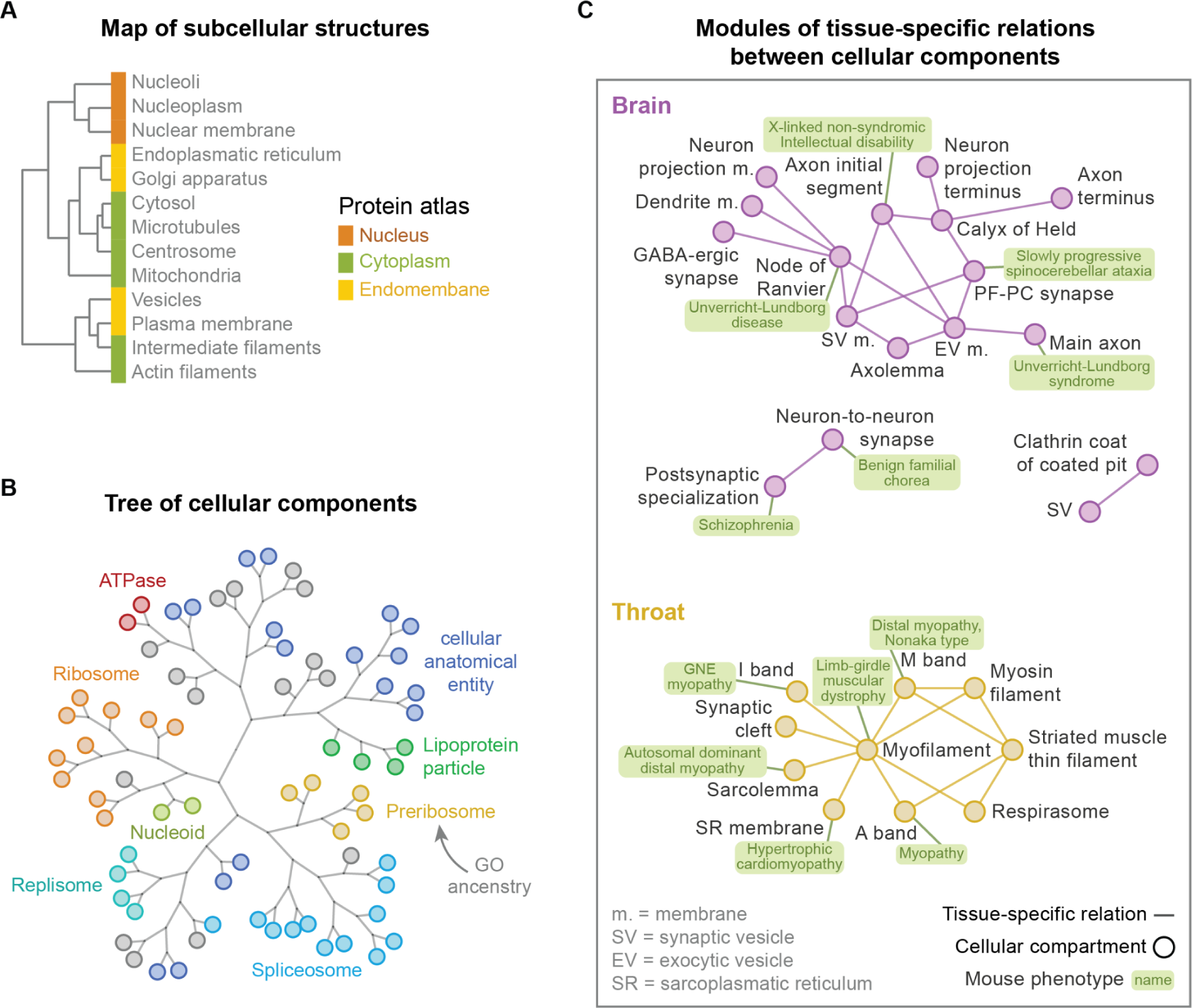
Mapping the organization of subcellular structures (Related to Fig. 3). Using our association atlas, we sought to explore the structural organization of the human cell by systematically mapping the relations between multi-protein structures such as organelles and protein complexes in a tissue-specific manner. Most ontologies such as GO have been built and manually curated by domain experts. However, manually curating relations between ontology terms at scale and in a consistent manner has proven to be difficult. Moreover, systematically mapping the relations between ontology terms is challenging as it requires some relational information to link terms with no overlapping genes. For example, pairs of protein sets such as GO cellular components share few proteins with an average Jaccard index of ∼0.008, illustrating that such gene sets are insufficient to link arbitrary pairs of ontology terms. However, as an example, one would expect that proteins that are annotated to be related with the nuclear membrane have higher association scores with proteins annotated with other nuclear-linked compartments compared to arbitrary proteins. By calculating the average association scores between pairs of proteins from different components for each tissue, we could thus build a tree of relationships that recovers the structural organization of the cell in a tissue-specific manner. We defined the relationship score between two ontology terms as the median association score between all pairs of proteins that are not shared between the terms (following Fig. 3). This methodology thus omits any similarity between terms that is simply due to the presence of common genes. **(a)** As a proof-of-concept, we sought to explore the relationships between subcellular structures as defined through cellular location data reported by the Protein Atlas (date = 23-06-17). We filtered for proteins that had an unique main location and whose reliability was ‘Enhanced’ or ‘Supported’. We then determined the relationship scores between all subcellular locations in all tissues. Finally, we averaged the relationship scores between subcellular locations across tissues, and then clustered the locations to reconstruct a map of subcellular structures. Dendrogram shows the subcellular structures as defined by cellular location of proteins from the Protein Atlas (complete-linkage clustering with the Manhattan distance). Subcellular structures are labeled as nuclear (orange), cytoplasm (green) or endomembrane (yellow) following annotation from the Protein Atlas. We found that the structures were organized based on both organellar similarity (i.e., nuclear, cytoplasmic and endomembrane) and spatial distribution (i.e., nucleus, cellular core and periphery). The relationship scores thus reveal a hierarchical organization of structures in the cell. **(b)** With the same approach, we used the relationship scores to map the hierarchical organization of GO cellular components. We computed the component-level association scores as the median association score between all pairs of proteins associated with a GO cellular component. We then selected the cellular components whose median association score exceeded 0.5 when averaged over the tissues. We then used the relationship scores between these 71 cellular components, relative to the median association score of each tissue, to compute the average relative relation between each pair of components. Dendrogram shows hierarchical organization of GO cellular components (complete linkage clustering with the Manhattan distance of the relationship matrix). Leaves are colored based on shared GO ancestry of the cellular components (i.e., components are colored based on shared first common ancestors in the go-basic dataset (date=27-07-2023). Specifically, following the branches from leaves to the root, each intermediate cluster in the dendrogram was labeled according to the first common ancestor of its leaves. Only ‘is_a’ relationships of cellular compartments in go-basic were considered (goatools)). We found that cellular components with common GO ancestry generally clustered together, except for a diverse group of cellular anatomical entities. The proposed methodology thus reconstructs the spatial organization of preserved cellular structures on several size-scales ranging from protein complexes to subcellular compartments. **(c)** Finally, we used the relationship scores across human tissues to map the tissue-specificity of cellular components. As an example, for each tissue, we filtered for pairs of components that consisted of at least 15 proteins and whose relationship score exceeded the 99.5-th percentile of all pairs of components. We then selected pairs of cellular components whose relationship scores were tissue-specific (z-scores > 3), revealing modules of high-confidence, tissue-specific relations between cellular components for the brain (purple) and throat (yellow). We annotated the cellular components that were enriched with genes from mouse phenotypes (green boxes - IMPC). Here, each component shows at most one phenotype and each phenotype is shown only once, for the component having the strongest enrichment (Fisher’s exact test, Benjamini-Hochberg (BH) adjusted p-values < 0.0001). The brain modules had tissue-specific connections involving synaptic components that were enriched with genes associated with brain disorders in mice. Similarly, for the throat, we found that the modules had tissue-specific connections of cellular components for muscle fiber and were enriched with genes associated with muscular disorders in mice. Overall, these analyses demonstrate the use of our association atlas for systematically scoring the relations between cellular structures in a tissue-specific manner.

**Figure S13.**
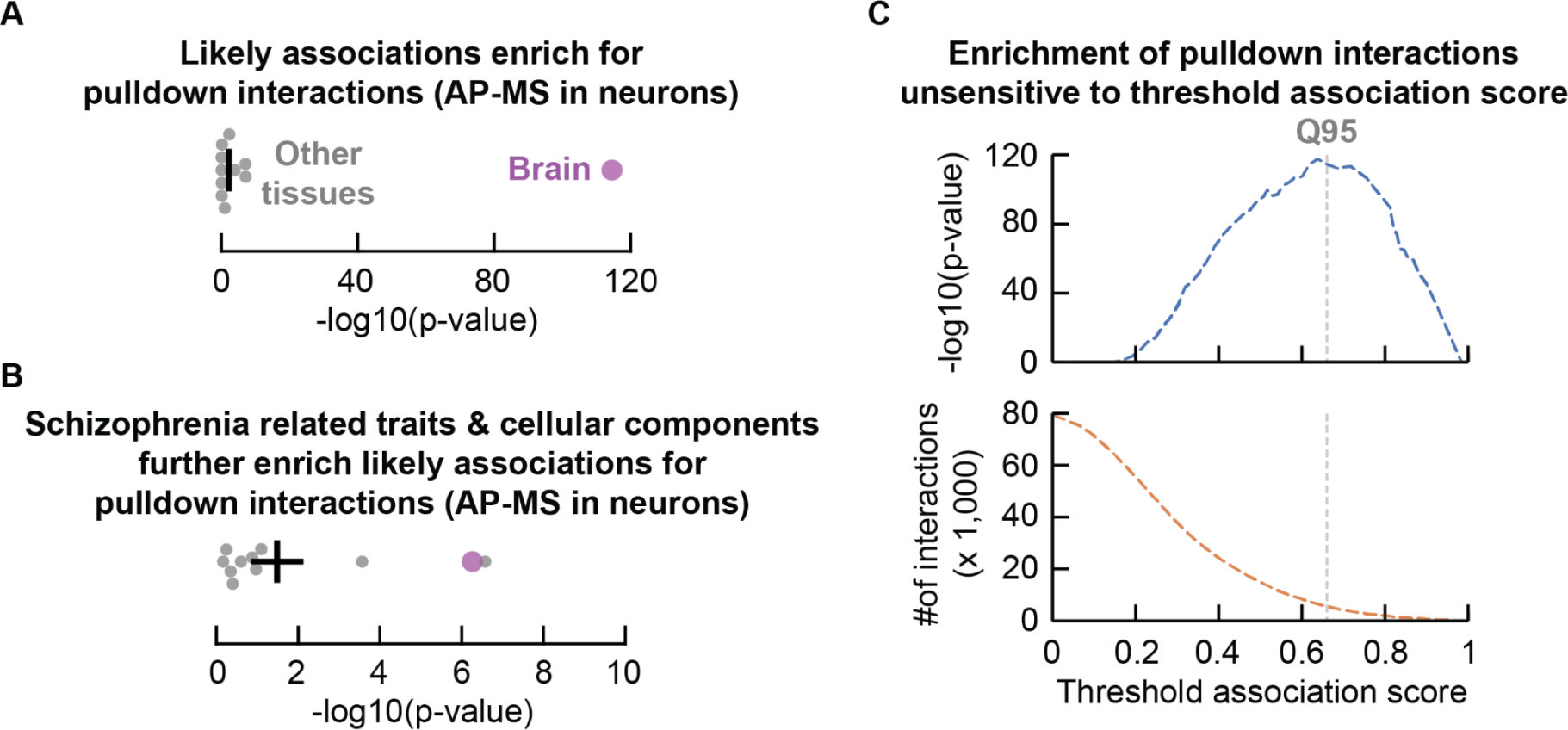
Schizophrenia-related brain associations enriched for interactions from AP-MS in neurons (Related to Fig. 4). As before, we filtered the brain interactions from pulldowns to baits that we would consider based on GWAS studies (L2G > 0.5), and we filtered the protein associations in each tissue to only include associations that we could have detected with the pulldown studies (i.e., interactions of the bait proteins). We then tested how well the association scores could recover the brain interactions from pulldown studies in a tissue-specific manner. **(a)** First, we tested whether the most likely associations of each tissue (association scores exceeding the 95-th percentile) were enriched for these pulldown interactions. Indeed, we found that these top-percentile associations were enriched for the brain-interactions from pulldowns, especially for the brain (log10 p-value 114.6) compared to the other tissues (2.2 p/m 0.9). **(b)** Next, we tested whether we could filter these most likely associations to further enrich the pulldown interactions. To do so, we selected the associations of protein pairs that involved one schizophrenia starting gene (L2G > 0.5) and one protein from the top 15 traits and cellular components that had the strongest tissue-specific relation to schizophrenia in each tissue (Methods). Here, we found that these schizophrenia-prioritized protein pairs were further enriched for the pulldown interactions compared to the other likely interactions, with the brain (-log10 p-value 6.3) outperforming most other tissues (1.5 p/m 0.6 - except for the lung (6.6)). We could thus use the likely associations of protein pairs involving schizophrenia genes and genes from related traits and cellular components to enrich for interactions found for schizophrenia genes with pulldowns in brain cells. **(c)** Finally, we confirmed that the enrichment of pulldown interactions amongst the likely schizophrenia-prioritized associations was not sensitive to the choice of the threshold association score (i.e., the log10 p-value exceeded 45 for all threshold association scores in the range 0.5 (all likely interactions; ∼85-th percentile) to 0.9 (∼99.8-th percentile)).

**Figure S14.**
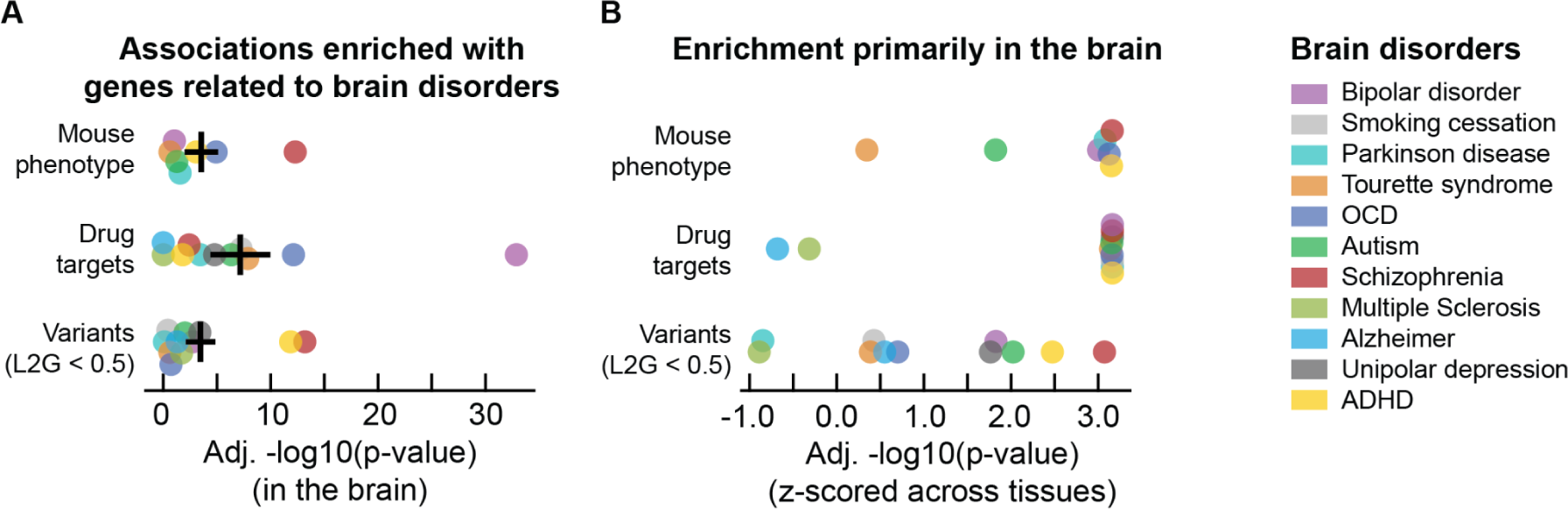
Proposed methodology for constructing networks of disease-related brain associations enriches for disease genes (Related to Fig. 4). We selected brain-disorders from the GWAS traits that were also a mouse phenotype (IMPC) or had drug targets (ChEMBL) and whose trait-level association scores were elevated in the brain compared to the other tissues (z-score > 0; Fig. 3). Following the methodology identical to selecting the potential schizophrenia-related interactions (Fig. 4a-b), we created tissue-specific networks of associations for each of the resulting 11 brain disorders. In short, we started by taking the genes associated with each disorder through GWAS studies (“starting genes” – L2G scores > 0.5) and computed the top 15 traits and cellular components that had the strongest tissue-specific relation to the respective disorder in each tissue. We then considered as potential interactions all protein pairs that had one starting gene for a brain-disorder and one related gene for that same disorder, respectively (Methods). Next, for each tissue, we filtered these potential interactions for being in the 95-th percentile of the tissue-scores, leading to tissue-specific networks of associations for each brain-disorder. **(a)** After removing the starting genes for each disorder from the respective brain-network, the remaining genes were enriched for genes associated to the respective disorders through mouse phenotypes (log10-p-value ∼3.5 p/m 1.5 – Fisher’s exact test), drug targets (∼7.2 p/m 2.8 – ChEMBL clinical stage II and above), and variants with weak evidence supporting them as causal for the respective disorder (∼3.5 p/m 1.4 – L2G scores < 0.5). Shown are the BH-adjusted log10-p-values for each disorder. **(b)** For most disorders (except Alzheimer, Parkinson, and multiple sclerosis), this enrichment was elevated for the brain compared to the other tissues, as demonstrated by the z-scored adjusted log10-p-values for the brain (z-score > 0). This enrichment was especially striking for autism, bipolar disorder, schizophrenia and ADHD (z-score > 1.5) for all of the gene-disease associations. Together, these observations suggest that the proposed methodology presents a systematic approach for prioritizing disease genes of tissue-specific traits.

## Supplementary Tables

Table S1 to Table S31 are available through the BioStudies repository for this paper https://www.ebi.ac.uk/biostudies/studies/S-BSST1423.

## References

1. Drew K, Wallingford JB, Marcotte EM. hu.MAP 2.0: integration of over 15,000 proteomic experiments builds a global compendium of human multiprotein assemblies. Mol Syst Biol. 2021;17: e10016.

2. Tarassov K, Messier V, Landry CR, Radinovic S, Serna Molina MM, Shames I, et al. An in vivo map of the yeast protein interactome. Science. 2008;320: 1465–1470.

3. Luck K, Kim D-K, Lambourne L, Spirohn K, Begg BE, Bian W, et al. A reference map of the human binary protein interactome. Nature. 2020;580: 402–408.

4. Huttlin EL, Bruckner RJ, Navarrete-Perea J, Cannon JR, Baltier K, Gebreab F, et al. Dual proteome-scale networks reveal cell-specific remodeling of the human interactome. Cell. 2021. doi:10.1016/j.cell.2021.04.011

5. Hein MY, Hubner NC, Poser I, Cox J, Nagaraj N, Toyoda Y, et al. A human interactome in three quantitative dimensions organized by stoichiometries and abundances. Cell. 2015;163: 712–723.

6. Skinnider MA, Scott NE, Prudova A, Kerr CH, Stoynov N, Stacey RG, et al. An atlas of protein-protein interactions across mouse tissues. Cell. 2021;184: 4073–4089.e17.

7. Orre LM, Vesterlund M, Pan Y, Arslan T, Zhu Y, Fernandez Woodbridge A, et al. SubCellBarCode: Proteome-wide Mapping of Protein Localization and Relocalization. Mol Cell. 2019;73: 166–182.e7.

8. Havugimana PC, Hart GT, Nepusz T, Yang H, Turinsky AL, Li Z, et al. A census of human soluble protein complexes. Cell. 2012;150: 1068–1081.

9. Orchard S, Ammari M, Aranda B, Breuza L, Briganti L, Broackes-Carter F, et al. The MIntAct project--IntAct as a common curation platform for 11 molecular interaction databases. Nucleic Acids Res. 2014;42: D358–63.

10. Oughtred R, Rust J, Chang C, Breitkreutz B-J, Stark C, Willems A, et al. The BioGRID database: A comprehensive biomedical resource of curated protein, genetic, and chemical interactions. Protein Sci. 2021;30: 187–200.

11. Szklarczyk D, Gable AL, Nastou KC, Lyon D, Kirsch R, Pyysalo S, et al. The STRING database in 2021: customizable protein-protein networks, and functional characterization of user-uploaded gene/measurement sets. Nucleic Acids Res. 2021;49: D605–D612.

12. Uhlén M, Fagerberg L, Hallström BM, Lindskog C, Oksvold P, Mardinoglu A, et al. Proteomics. Tissue-based map of the human proteome. Science. 2015;347: 1260419.

13. Pierson E, GTEx Consortium, Koller D, Battle A, Mostafavi S, Ardlie KG, et al. Sharing and Specificity of Co-expression Networks across 35 Human Tissues. PLoS Comput Biol. 2015;11: e1004220.

14. Greene CS, Krishnan A, Wong AK, Ricciotti E, Zelaya RA, Himmelstein DS, et al. Understanding multicellular function and disease with human tissue-specific networks. Nat Genet. 2015;47: 569–576.

15. Sousa A, Gonçalves E, Mirauta B, Ochoa D, Stegle O, Beltrao P. Multi-omics Characterization of Interaction-mediated Control of Human Protein Abundance levels. Mol Cell Proteomics. 2019;18: S114–S125.

16. Ryan CJ, Kennedy S, Bajrami I, Matallanas D, Lord CJ. A Compendium of Co-regulated Protein Complexes in Breast Cancer Reveals Collateral Loss Events. Cell Syst. 2017;5: 399–409.e5.

17. Lapek JD Jr, Greninger P, Morris R, Amzallag A, Pruteanu-Malinici I, Benes CH, et al. Detection of dysregulated protein-association networks by high-throughput proteomics predicts cancer vulnerabilities. Nat Biotechnol. 2017;35: 983–989.

18. Wang J, Ma Z, Carr SA, Mertins P, Zhang H, Zhang Z, et al. Proteome Profiling Outperforms Transcriptome Profiling for Coexpression Based Gene Function Prediction. Mol Cell Proteomics. 2017;16: 121–134.

19. Buljan M, Banaei-Esfahani A, Blattmann P, Meier-Abt F, Shao W, Vitek O, et al. A computational framework for the inference of protein complex remodeling from whole-proteome measurements. Nat Methods. 2023;20: 1523–1529.

20. Roumeliotis TI, Williams SP, Gonçalves E, Alsinet C, Del Castillo Velasco-Herrera M, Aben N, et al. Genomic Determinants of Protein Abundance Variation in Colorectal Cancer Cells. Cell Rep. 2017;20: 2201–2214.

21. Kustatscher G, Grabowski P, Schrader TA, Passmore JB, Schrader M, Rappsilber J. Co-regulation map of the human proteome enables identification of protein functions. Nat Biotechnol. 2019;37: 1361–1371.

22. Taggart JC, Zauber H, Selbach M, Li G-W, McShane E. Keeping the Proportions of Protein Complex Components in Check. Cell Syst. 2020;10: 125–132.

23. Juszkiewicz S, Hegde RS. Quality Control of Orphaned Proteins. Mol Cell. 2018;71: 443–457.

24. Gonçalves E, Fragoulis A, Garcia-Alonso L, Cramer T, Saez-Rodriguez J, Beltrao P. Widespread Post-transcriptional Attenuation of Genomic Copy-Number Variation in Cancer. Cell Syst. 2017;5: 386–398.e4.

25. McShane E, Sin C, Zauber H, Wells JN, Donnelly N, Wang X, et al. Kinetic Analysis of Protein Stability Reveals Age-Dependent Degradation. Cell. 2016;167: 803–815.e21.

26. Xu N, Yao Z, Shang G, Ye D, Wang H, Zhang H, et al. Integrated proteogenomic characterization of urothelial carcinoma of the bladder. J Hematol Oncol. 2022;15: 76.

27. Herbst SA, Vesterlund M, Helmboldt AJ, Jafari R, Siavelis I, Stahl M, et al. Proteogenomics refines the molecular classification of chronic lymphocytic leukemia. Nat Commun. 2022;13: 6226.

28. Jayavelu AK, Wolf S, Buettner F, Alexe G, Häupl B, Comoglio F, et al. The proteogenomic subtypes of acute myeloid leukemia. Cancer Cell. 2022;40: 301–317.e12.

29. Kramer MH, Zhang Q, Sprung R, Day RB, Erdmann-Gilmore P, Li Y, et al. Proteomic and phosphoproteomic landscapes of acute myeloid leukemia. Blood. 2022;140: 1533–1548.

30. Stratmann S, Vesterlund M, Umer HM, Eshtad S, Skaftason A, Herlin MK, et al. Proteogenomic analysis of acute myeloid leukemia associates relapsed disease with reprogrammed energy metabolism both in adults and children. Leukemia. 2023;37: 550–559.

31. Yang M, Vesterlund M, Siavelis I, Moura-Castro LH, Castor A, Fioretos T, et al. Proteogenomics and Hi-C reveal transcriptional dysregulation in high hyperdiploid childhood acute lymphoblastic leukemia. Nat Commun. 2019;10: 1519.

32. Archer TC, Ehrenberger T, Mundt F, Gold MP, Krug K, Mah CK, et al. Proteomics, Post-translational Modifications, and Integrative Analyses Reveal Molecular Heterogeneity within Medulloblastoma Subgroups. Cancer Cell. 2018;34: 396–410.e8.

33. Oh S, Yeom J, Cho HJ, Kim J-H, Yoon S-J, Kim H, et al. Integrated pharmaco-proteogenomics defines two subgroups in isocitrate dehydrogenase wild-type glioblastoma with prognostic and therapeutic opportunities. Nat Commun. 2020;11: 3288.

34. Petralia F, Tignor N, Reva B, Koptyra M, Chowdhury S, Rykunov D, et al. Integrated Proteogenomic Characterization across Major Histological Types of Pediatric Brain Cancer. Cell. 2020;183: 1962–1985.e31.

35. Wang L-B, Karpova A, Gritsenko MA, Kyle JE, Cao S, Li Y, et al. Proteogenomic and metabolomic characterization of human glioblastoma. Cancer Cell. 2021;39: 509–528.e20.

36. Yanovich-Arad G, Ofek P, Yeini E, Mardamshina M, Danilevsky A, Shomron N, et al. Proteogenomics of glioblastoma associates molecular patterns with survival. Cell Rep. 2021;34: 108787.

37. Zhang F, Zhang Q, Zhu J, Yao B, Ma C, Qiao N, et al. Integrated proteogenomic characterization across major histological types of pituitary neuroendocrine tumors. Cell Res. 2022;32: 1047–1067.

38. Anurag M, Jaehnig EJ, Krug K, Lei JT, Bergstrom EJ, Kim B-J, et al. Proteogenomic Markers of Chemotherapy Resistance and Response in Triple-Negative Breast Cancer. Cancer Discov. 2022;12: 2586–2605.

39. Asleh K, Negri GL, Spencer Miko SE, Colborne S, Hughes CS, Wang XQ, et al. Proteomic analysis of archival breast cancer clinical specimens identifies biological subtypes with distinct survival outcomes. Nat Commun. 2022;13: 896.

40. Gong T-Q, Jiang Y-Z, Shao C, Peng W-T, Liu M-W, Li D-Q, et al. Proteome-centric cross-omics characterization and integrated network analyses of triple-negative breast cancer. Cell Rep. 2022;38: 110460.

41. Johansson HJ, Socciarelli F, Vacanti NM, Haugen MH, Zhu Y, Siavelis I, et al. Breast cancer quantitative proteome and proteogenomic landscape. Nat Commun. 2019;10: 1600.

42. Krug K, Jaehnig EJ, Satpathy S, Blumenberg L, Karpova A, Anurag M, et al. Proteogenomic Landscape of Breast Cancer Tumorigenesis and Targeted Therapy. Cell. 2020;183: 1436–1456.e31.

43. Mertins P, Mani DR, Ruggles KV, Gillette MA, Clauser KR, Wang P, et al. Proteogenomics connects somatic mutations to signalling in breast cancer. Nature. 2016;534: 55–62.

44. Li C, Sun Y-D, Yu G-Y, Cui J-R, Lou Z, Zhang H, et al. Integrated Omics of Metastatic Colorectal Cancer. Cancer Cell. 2020;38: 734–747.e9.

45. Vasaikar S, Huang C, Wang X, Petyuk VA, Savage SR, Wen B, et al. Proteogenomic Analysis of Human Colon Cancer Reveals New Therapeutic Opportunities. Cell. 2019;177: 1035–1049.e19.

46. Wang J, Mouradov D, Wang X, Jorissen RN, Chambers MC, Zimmerman LJ, et al. Colorectal Cancer Cell Line Proteomes Are Representative of Primary Tumors and Predict Drug Sensitivity. Gastroenterology. 2017;153: 1082–1095.

47. Clark DJ, Dhanasekaran SM, Petralia F, Pan J, Song X, Hu Y, et al. Integrated Proteogenomic Characterization of Clear Cell Renal Cell Carcinoma. Cell. 2019;179: 964–983.e31.

48. Li Y, Lih T-SM, Dhanasekaran SM, Mannan R, Chen L, Cieslik M, et al. Histopathologic and proteogenomic heterogeneity reveals features of clear cell renal cell carcinoma aggressiveness. Cancer Cell. 2023;41: 139–163.e17.

49. Qu Y, Feng J, Wu X, Bai L, Xu W, Zhu L, et al. A proteogenomic analysis of clear cell renal cell carcinoma in a Chinese population. Nat Commun. 2022;13: 2052.

50. Qu Y, Wu X, Anwaier A, Feng J, Xu W, Pei X, et al. Proteogenomic characterization of MiT family translocation renal cell carcinoma. Nat Commun. 2022;13: 7494.

51. Dong L, Lu D, Chen R, Lin Y, Zhu H, Zhang Z, et al. Proteogenomic characterization identifies clinically relevant subgroups of intrahepatic cholangiocarcinoma. Cancer Cell. 2022;40: 70–87.e15.

52. Gao Q, Zhu H, Dong L, Shi W, Chen R, Song Z, et al. Integrated Proteogenomic Characterization of HBV-Related Hepatocellular Carcinoma. Cell. 2019;179: 561–577.e22.

53. Gu Y, Guo Y, Gao N, Fang Y, Xu C, Hu G, et al. The proteomic characterization of the peritumor microenvironment in human hepatocellular carcinoma. Oncogene. 2022;41: 2480–2491.

54. Jiang Y, Sun A, Zhao Y, Ying W, Sun H, Yang X, et al. Proteomics identifies new therapeutic targets of early-stage hepatocellular carcinoma. Nature. 2019;567: 257–261.

55. Ng CKY, Dazert E, Boldanova T, Coto-Llerena M, Nuciforo S, Ercan C, et al. Integrative proteogenomic characterization of hepatocellular carcinoma across etiologies and stages. Nat Commun. 2022;13: 2436.

56. Chen Y-J, Roumeliotis TI, Chang Y-H, Chen C-T, Han C-L, Lin M-H, et al. Proteogenomics of Non-smoking Lung Cancer in East Asia Delineates Molecular Signatures of Pathogenesis and Progression. Cell. 2020;182: 226–244.e17.

57. Gillette MA, Satpathy S, Cao S, Dhanasekaran SM, Vasaikar SV, Krug K, et al. Proteogenomic Characterization Reveals Therapeutic Vulnerabilities in Lung Adenocarcinoma. Cell. 2020;182: 200–225.e35.

58. Lehtiö J, Arslan T, Siavelis I, Pan Y, Socciarelli F, Berkovska O, et al. Proteogenomics of non-small cell lung cancer reveals molecular subtypes associated with specific therapeutic targets and immune evasion mechanisms. Nat Cancer. 2021;2: 1224–1242.

59. Satpathy S, Krug K, Jean Beltran PM, Savage SR, Petralia F, Kumar-Sinha C, et al. A proteogenomic portrait of lung squamous cell carcinoma. Cell. 2021;184: 4348–4371.e40.

60. Soltis AR, Bateman NW, Liu J, Nguyen T, Franks TJ, Zhang X, et al. Proteogenomic analysis of lung adenocarcinoma reveals tumor heterogeneity, survival determinants, and therapeutically relevant pathways. Cell Rep Med. 2022;3: 100819.

61. Stewart PA, Welsh EA, Slebos RJC, Fang B, Izumi V, Chambers M, et al. Proteogenomic landscape of squamous cell lung cancer. Nat Commun. 2019;10: 3578.

62. Xu J-Y, Zhang C, Wang X, Zhai L, Ma Y, Mao Y, et al. Integrative Proteomic Characterization of Human Lung Adenocarcinoma. Cell. 2020;182: 245–261.e17.

63. McDermott JE, Arshad OA, Petyuk VA, Fu Y, Gritsenko MA, Clauss TR, et al. Proteogenomic Characterization of Ovarian HGSC Implicates Mitotic Kinases, Replication Stress in Observed Chromosomal Instability. Cell Rep Med. 2020;1. doi:10.1016/j.xcrm.2020.100004

64. Zhang H, Liu T, Zhang Z, Payne SH, Zhang B, McDermott JE, et al. Integrated Proteogenomic Characterization of Human High-Grade Serous Ovarian Cancer. Cell. 2016;166: 755–765.

65. Cao L, Huang C, Cui Zhou D, Hu Y, Lih TM, Savage SR, et al. Proteogenomic characterization of pancreatic ductal adenocarcinoma. Cell. 2021;184: 5031–5052.e26.

66. Hyeon DY, Nam D, Han Y, Kim DK, Kim G, Kim D, et al. Proteogenomic landscape of human pancreatic ductal adenocarcinoma in an Asian population reveals tumor cell-enriched and immune-rich subtypes. Nat Cancer. 2023;4: 290–307.

67. Sinha A, Huang V, Livingstone J, Wang J, Fox NS, Kurganovs N, et al. The Proteogenomic Landscape of Curable Prostate Cancer. Cancer Cell. 2019;35: 414–427.e6.

68. Fan Y, Bai B, Liang Y, Ren Y, Liu Y, Zhou F, et al. Proteomic Profiling of Gastric Signet Ring Cell Carcinoma Tissues Reveals Characteristic Changes of the Complement Cascade Pathway. Mol Cell Proteomics. 2021;20: 100068.

69. Ge S, Xia X, Ding C, Zhen B, Zhou Q, Feng J, et al. A proteomic landscape of diffuse-type gastric cancer. Nat Commun. 2018;9: 1012.

70. Li Y, Xu C, Wang B, Xu F, Ma F, Qu Y, et al. Proteomic characterization of gastric cancer response to chemotherapy and targeted therapy reveals new therapeutic strategies. Nat Commun. 2022;13: 5723.

71. Huang C, Chen L, Savage SR, Eguez RV, Dou Y, Li Y, et al. Proteogenomic insights into the biology and treatment of HPV-negative head and neck squamous cell carcinoma. Cancer Cell. 2021;39: 361–379.e16.

72. Li S, Yuan L, Xu Z-Y, Xu J-L, Chen G-P, Guan X, et al. Integrative proteomic characterization of adenocarcinoma of esophagogastric junction. Nat Commun. 2023;14: 778.

73. Liu W, Xie L, He Y-H, Wu Z-Y, Liu L-X, Bai X-F, et al. Large-scale and high-resolution mass spectrometry-based proteomics profiling defines molecular subtypes of esophageal cancer for therapeutic targeting. Nat Commun. 2021;12: 4961.

74. Bateman NW, Tarney CM, Abulez T, Soltis AR, Zhou M, Conrads K, et al. Proteogenomic landscape of uterine leiomyomas from hereditary leiomyomatosis and renal cell cancer patients. Sci Rep. 2021;11: 9371.

75. Dou Y, Kawaler EA, Cui Zhou D, Gritsenko MA, Huang C, Blumenberg L, et al. Proteogenomic Characterization of Endometrial Carcinoma. Cell. 2020;180: 729–748.e26.

76. Ruepp A, Waegele B, Lechner M, Brauner B, Dunger-Kaltenbach I, Fobo G, et al. CORUM: the comprehensive resource of mammalian protein complexes--2009. Nucleic Acids Res. 2010;38: D497–501.

77. Havugimana PC, Goel RK, Phanse S, Youssef A, Padhorny D, Kotelnikov S, et al. Scalable multiplex co-fractionation/mass spectrometry platform for accelerated protein interactome discovery. Nat Commun. 2022;13: 4043.

78. Lo Surdo P, Iannuccelli M, Contino S, Castagnoli L, Licata L, Cesareni G, et al. SIGNOR 3.0, the SIGnaling network open resource 3.0: 2022 update. Nucleic Acids Res. 2023;51: D631–D637.

79. Milacic M, Beavers D, Conley P, Gong C, Gillespie M, Griss J, et al. The Reactome Pathway Knowledgebase 2024. Nucleic Acids Res. 2024;52: D672–D678.

80. Szklarczyk D, Gable AL, Lyon D, Junge A, Wyder S, Huerta-Cepas J, et al. STRING v11: protein-protein association networks with increased coverage, supporting functional discovery in genome-wide experimental datasets. Nucleic Acids Res. 2019;47: D607–D613.

81. Romanov N, Kuhn M, Aebersold R, Ori A, Beck M, Bork P. Disentangling Genetic and Environmental Effects on the Proteotypes of Individuals. Cell. 2019;177: 1308–1318.e10.

82. Guardia CM, De Pace R, Mattera R, Bonifacino JS. Neuronal functions of adaptor complexes involved in protein sorting. Curr Opin Neurobiol. 2018;51: 103–110.

83. Koopmans F, van Nierop P, Andres-Alonso M, Byrnes A, Cijsouw T, Coba MP, et al. SynGO: An Evidence-Based, Expert-Curated Knowledge Base for the Synapse. Neuron. 2019;103: 217–234.e4.

84. Huang L-H, Deepak P, Ciorba MA, Mittendorfer B, Patterson BW, Randolph GJ. Postprandial Chylomicron Output and Transport Through Intestinal Lymphatics Are Not Impaired in Active Crohn’s Disease. Gastroenterology. 2020;159: 1955–1957.e2.

85. Ghoshal S, Witta J, Zhong J, de Villiers W, Eckhardt E. Chylomicrons promote intestinal absorption of lipopolysaccharides. J Lipid Res. 2009;50: 90–97.

86. Kopec AK, Luyendyk JP. Role of Fibrin(ogen) in Progression of Liver Disease: Guilt by Association? Semin Thromb Hemost. 2016;42: 397–407.

87. Northup PG, Caldwell SH. Coagulation in liver disease: a guide for the clinician. Clin Gastroenterol Hepatol. 2013;11: 1064–1074.

88. Ashburner M, Ball CA, Blake JA, Botstein D, Butler H, Cherry JM, et al. Gene ontology: tool for the unification of biology. The Gene Ontology Consortium. Nat Genet. 2000;25: 25–29.

89. Ghoussaini M, Mountjoy E, Carmona M, Peat G, Schmidt EM, Hercules A, et al. Open Targets Genetics: systematic identification of trait-associated genes using large-scale genetics and functional genomics. Nucleic Acids Res. 2021;49: D1311–D1320.

90. Mountjoy E, Schmidt EM, Carmona M, Schwartzentruber J, Peat G, Miranda A, et al. An open approach to systematically prioritize causal variants and genes at all published human GWAS trait-associated loci. Nat Genet. 2021;53: 1527–1533.

91. Yan J, Deng H, Wang Y, Wang X, Fan T, Li S, et al. The Prevalence and Comorbidity of Tic Disorders and Obsessive-Compulsive Disorder in Chinese School Students Aged 6-16: A National Survey. Brain Sci. 2022;12. doi:10.3390/brainsci12050650

92. Kumar A, Trescher W, Byler D. Tourette Syndrome and Comorbid Neuropsychiatric Conditions. Curr Dev Disord Rep. 2016;3: 217–221.

93. Davies M, Nowotka M, Papadatos G, Dedman N, Gaulton A, Atkinson F, et al. ChEMBL web services: streamlining access to drug discovery data and utilities. Nucleic Acids Res. 2015;43: W612–20.

94. Dickinson ME, Flenniken AM, Ji X, Teboul L, Wong MD, White JK, et al. High-throughput discovery of novel developmental phenotypes. Nature. 2016;537: 508–514.

95. Sadler MC, Auwerx C, Deelen P, Kutalik Z. Multi-layered genetic approaches to identify approved drug targets. Cell Genom. 2023;3: 100341.

96. Barrio-Hernandez I, Schwartzentruber J, Shrivastava A, Del-Toro N, Gonzalez A, Zhang Q, et al. Network expansion of genetic associations defines a pleiotropy map of human cell biology. Nat Genet. 2023;55: 389–398.

97. Lee I, Blom UM, Wang PI, Shim JE, Marcotte EM. Prioritizing candidate disease genes by network-based boosting of genome-wide association data. Genome Res. 2011;21: 1109–1121.

98. Hsu Y-HH, Pintacuda G, Liu R, Nacu E, Kim A, Tsafou K, et al. Using brain cell-type-specific protein interactomes to interpret neurodevelopmental genetic signals in schizophrenia. iScience. 2023;26: 106701.

99. Pintacuda G, Hsu Y-HH, Tsafou K, Li KW, Martín JM, Riseman J, et al. Protein interaction studies in human induced neurons indicate convergent biology underlying autism spectrum disorders. Cell Genom. 2023;3: 100250.

100. O’Neill AC, Uzbas F, Antognolli G, Merino F, Draganova K, Jäck A, et al. Spatial centrosome proteome of human neural cells uncovers disease-relevant heterogeneity. Science. 2022;376: eabf9088.

101. Drummond E, Pires G, MacMurray C, Askenazi M, Nayak S, Bourdon M, et al. Phosphorylated tau interactome in the human Alzheimer’s disease brain. Brain. 2020;143: 2803–2817.

102. Tracy TE, Madero-Pérez J, Swaney DL, Chang TS, Moritz M, Konrad C, et al. Tau interactome maps synaptic and mitochondrial processes associated with neurodegeneration. Cell. 2022;185: 712–728.e14.

103. Runge K, Balla A, Fiebich BL, Maier SJ, von Zedtwitz K, Nickel K, et al. Neurodegeneration Markers in the Cerebrospinal Fluid of 100 Patients with Schizophrenia Spectrum Disorder. Schizophr Bull. 2023;49: 464–473.

104. Lewis AS, Schwartz E, Chan CS, Noam Y, Shin M, Wadman WJ, et al. Alternatively spliced isoforms of TRIP8b differentially control h channel trafficking and function. J Neurosci. 2009;29: 6250–6265.

105. Santoro B, Hu L, Liu H, Saponaro A, Pian P, Piskorowski RA, et al. TRIP8b regulates HCN1 channel trafficking and gating through two distinct C-terminal interaction sites. J Neurosci. 2011;31: 4074–4086.

106. Popova NV, Plotnikov AN, Ziganshin RK, Deyev IE, Petrenko AG. Analysis of proteins interacting with TRIP8b adapter. Biochemistry. 2008;73: 644–651.

107. van Oostrum M, Blok TM, Giandomenico SL, Tom Dieck S, Tushev G, Fürst N, et al. The proteomic landscape of synaptic diversity across brain regions and cell types. Cell. 2023;186: 5411–5427.e23.

108. Roth FC, Hu H. An axon-specific expression of HCN channels catalyzes fast action potential signaling in GABAergic interneurons. Nat Commun. 2020;11: 2248.

109. Cai W, Liu S-S, Li B-M, Zhang X-H. Presynaptic HCN channels constrain GABAergic synaptic transmission in pyramidal cells of the medial prefrontal cortex. Biol Open. 2022;11. doi:10.1242/bio.058840

110. Kaar SJ, Angelescu I, Marques TR, Howes OD. Pre-frontal parvalbumin interneurons in schizophrenia: a meta-analysis of post-mortem studies. J Neural Transm. 2019;126: 1637–1651.

111. Arime Y, Saitoh Y, Ishikawa M, Kamiyoshihara C, Uchida Y, Fujii K, et al. Activation of prefrontal parvalbumin interneurons ameliorates working memory deficit even under clinically comparable antipsychotic treatment in a mouse model of schizophrenia. Neuropsychopharmacology. 2024;49: 720–730.

112. Hashimoto T, Volk DW, Eggan SM, Mirnics K, Pierri JN, Sun Z, et al. Gene expression deficits in a subclass of GABA neurons in the prefrontal cortex of subjects with schizophrenia. J Neurosci. 2003;23: 6315–6326.

113. Lewis DA, Hashimoto T, Volk DW. Cortical inhibitory neurons and schizophrenia. Nat Rev Neurosci. 2005;6: 312–324.

114. Bitanihirwe BKY, Lim MP, Kelley JF, Kaneko T, Woo TUW. Glutamatergic deficits and parvalbumin-containing inhibitory neurons in the prefrontal cortex in schizophrenia. BMC Psychiatry. 2009;9: 71.

115. Burke DF, Bryant P, Barrio-Hernandez I, Memon D, Pozzati G, Shenoy A, et al. Towards a structurally resolved human protein interaction network. Nat Struct Mol Biol. 2023;30: 216–225.

116. Madeira F, Tinti M, Murugesan G, Berrett E, Stafford M, Toth R, et al. 14-3-3-Pred: improved methods to predict 14-3-3-binding phosphopeptides. Bioinformatics. 2015;31: 2276–2283.

117. Lankford C, Houtman J, Baker SA. Identification of HCN1 as a 14-3-3 client. PLoS One. 2022;17: e0268335.

118. Gonzalez-Lozano MA, Koopmans F, Sullivan PF, Protze J, Krause G, Verhage M, et al. Stitching the synapse: Cross-linking mass spectrometry into resolving synaptic protein interactions. Sci Adv. 2020;6: eaax5783.

119. Moon HM, Hippenmeyer S, Luo L, Wynshaw-Boris A. LIS1 determines cleavage plane positioning by regulating actomyosin-mediated cell membrane contractility. Elife. 2020;9. doi:10.7554/eLife.51512

120. Skinnider MA, Foster LJ. Meta-analysis defines principles for the design and analysis of co-fractionation mass spectrometry experiments. Nat Methods. 2021;18: 806–815.

121. Frommelt F, Fossati A, Uliana F, Wendt F, Xue P, Heusel M, et al. DIP-MS: ultra-deep interaction proteomics for the deconvolution of protein complexes. Nat Methods. 2024;21: 635–647.

122. Liu Y, Mi Y, Mueller T, Kreibich S, Williams EG, Van Drogen A, et al. Multi-omic measurements of heterogeneity in HeLa cells across laboratories. Nat Biotechnol. 2019;37: 314–322.

123. Fossati A, Frommelt F, Uliana F, Martelli C, Vizovisek M, Gillet L, et al. System-Wide Profiling of Protein Complexes Via Size Exclusion Chromatography–Mass SpectrometryMass spectrometry (SEC–MS). In: Carrera M, Mateos J, editors. Shotgun Proteomics: Methods and Protocols. New York, NY: Springer US; 2021. pp. 269–294.

124. Bludau I, Heusel M, Frank M, Rosenberger G, Hafen R, Banaei-Esfahani A, et al. Complex-centric proteome profiling by SEC-SWATH-MS for the parallel detection of hundreds of protein complexes. Nat Protoc. 2020;15: 2341–2386.

125. Wildschut MHE, Mena J, Dördelmann C, van Oostrum M, Hale BD, Settelmeier J, et al. Proteogenetic drug response profiling elucidates targetable vulnerabilities of myelofibrosis. Nat Commun. 2023;14: 6414.

126. Muntel J, Kirkpatrick J, Bruderer R, Huang T, Vitek O, Ori A, et al. Comparison of Protein Quantification in a Complex Background by DIA and TMT Workflows with Fixed Instrument Time. J Proteome Res. 2019;18: 1340–1351.

127. Deutsch EW, Bandeira N, Perez-Riverol Y, Sharma V, Carver JJ, Mendoza L, et al. The ProteomeXchange consortium at 10 years: 2023 update. Nucleic Acids Res. 2023;51: D1539–D1548.

128. Perez-Riverol Y, Bai J, Bandla C, García-Seisdedos D, Hewapathirana S, Kamatchinathan S, et al. The PRIDE database resources in 2022: a hub for mass spectrometry-based proteomics evidences. Nucleic Acids Res. 2022;50: D543–D552.

